# Spatially resolved multimodal hallmarks of response to neoadjuvant immunotherapies in the melanoma ecosystem in 2D and 3D

**DOI:** 10.64898/2026.09.12.751186

**Authors:** Zichao Liu, Xiaofei Song, Wei-Shen Chen, Shikhar Dhingra, Jiang He, Lizhi He, Chengyi Chen, Jodi A. Balasi, Zena Sayegh, Carlos M. Moran Segura, Neale Lopez-Blanco, Anthony Alleyne, Jonathan V Nguyen, Joseph O. Johnson, Douglas Marchion, Sean J. Yoder, Jane L. Messina, Vernon K. Sondak, Joseph Markowitz, Nicholas P. Reder, Patrick Hwu, James J. Mulé, Jeffrey H. Chuang, Pei-Ling Chen

**Author notes:** Senior author (Jeffrey H. Chuang, Pei-Ling Chen).

## Abstract

Neoadjuvant immunotherapy has transformed cancer treatment, yet the spatial molecular architecture governing response and resistance across distinct immune checkpoint blockade (ICB) regimens remains poorly defined. We assembled the largest neoadjuvant ICB (NICB) spatial multi-omics cohort to date, profiling over 112 million cells at single-cell resolution across three melanoma ICB regimens using MERFISH spatial transcriptomics, multiplexed immunofluorescence, and scRNA-sequencing. These analyses revealed the full multicellular spatial architecture of the NICB tumor microenvironment, including mature TLS with germinal centers, TCF7+ stem-like T cells, myeloid cells organized into spatially distinct cellular neighborhoods with unique intercellular signaling circuits, and CCL19/CCL21-expressing fibroblasts as a previously unrecognized stromal scaffold sustaining these immune hubs. We developed three purpose-built computational tools that together enabled comprehensive quantification of this microenvironment for the first time: SCIRA for whole-slide single-cell receptor-ligand quantification, GCSCAN for molecularly grounded TLS and germinal center structural delineation, and PathNet-TLS for automated TLS detection on H&E images. Applying these tools across the cohort, we defined the immune and stromal composition and cellular neighborhood organization distinguishing responders from non-responders. We also quantified cell-cell interactions and regimen-specific immune architectures, including a markedly stronger mature TLS/germinal center response with IPI-NIVO than NIVO-RELA. Importantly, GCSCAN-quantified TLS and germinal center density each stratified disease-free survival, with responders that lack germinal centers having an elevated risk of relapse. Open-top light-sheet imaging and CODA-based 3D reconstruction further uncovered interconnected germinal center-TLS tunnels invisible to standard 2D histopathology. These findings establish a discovery-to-tool paradigm linking single-cell tumor microenvironment interrogation to clinically deployable computational pathology for biomarker-driven NICB assessment across cancer types.

**GRAPHICAL ABSTRACT:** 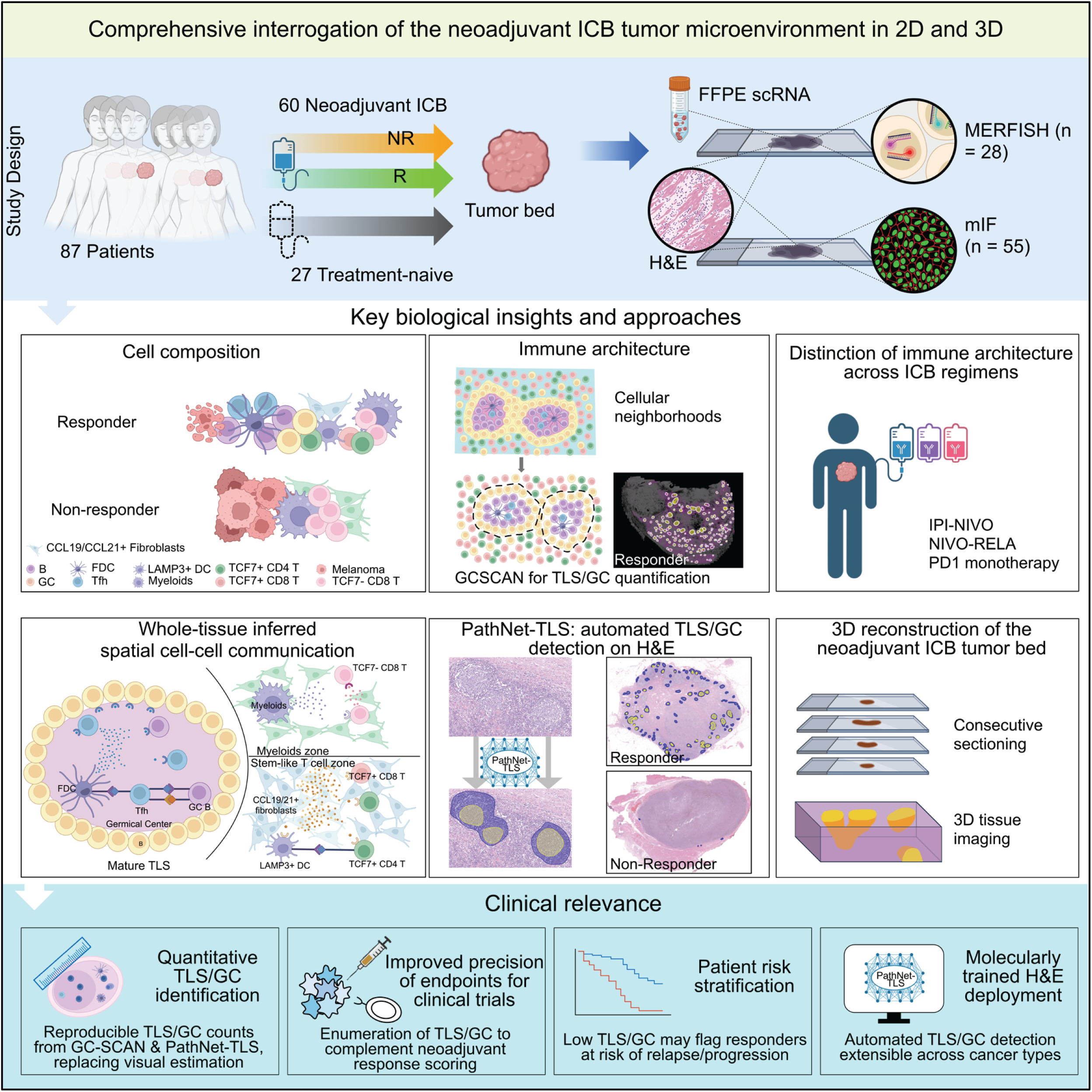

## INTRODUCTION

Immune checkpoint blockade (ICB) therapy represents a transformative advance in cancer treatment (1). Landmark neoadjuvant ICB clinical trials in melanoma have demonstrated decisively that administration of ICB prior to curative-intent surgery will become the new standard of care for resectable metastatic melanoma and, increasingly, across other cancer types (2–8). Beyond reducing tumor burden, neoadjuvant immunotherapy capitalizes on intact primary tumor antigens and relatively preserved T cell function to elicit a more robust and durable antitumor immune response (4,9,10). While neoadjuvant ICB clinical trials have demonstrated superior event-free survival compared to adjuvant therapy, a substantial proportion of patients fail to respond or develop acquired resistance (11,12), and reliable predictors of outcome remain lacking (13). A deeper understanding of how distinct ICB regimens modulate the tumor immune microenvironment is urgently needed to guide more effective treatment strategies and inform post-surgical personalized therapy decisions.

Melanoma has long been at the forefront of cancer immunotherapy, and the establishment of the International Neoadjuvant Melanoma Consortium (INMC) represented a landmark achievement in standardizing neoadjuvant pathologic response assessment criteria and aligning clinical trial design with correlative science (9,14). Despite these pioneering efforts, the current histologic response assessment (15,16) based on visual estimation of residual viable tumor, fibroinflammatory stroma, pigmented macrophages, and necrosis on H&E slides lacks precision and reproducibility, and a fundamental gap remains in our understanding of how spatial molecular interactions within the fibroinflammatory tumor microenvironment orchestrate antitumor responses across distinct ICB regimens. As the FDA has recognized pathologic response as a surrogate endpoint for accelerated approval across multiple cancer types (17,18), there is an urgent need for precise, scalable computational tools that can extract spatially resolved biomarkers from multiplexed assays and universally available H&E slides, both to improve response assessment and to elucidate the immunologic mechanisms underlying distinct neoadjuvant ICB regimens.

The spatially resolved molecular insights needed to inform such tools have only recently begun to emerge (19), even in melanoma, the cancer type at the forefront of ICB development. An initial groundbreaking scRNA-seq study identified CD8+ T cells with TCF7+ stem-like properties in lesions responding to adjuvant ICB (20), and subsequent multiplexed spatial profiling by imaging mass cytometry showed that B cell patches and follicles were enriched with TCF7+ naïve-like T cells (21). More recently, spatially organized immune hubs enriched for TCF7+ stem-like CD8 T cells, CCR7+LAMP3+ dendritic cells, and CCL19+ fibroblasts were identified in lung cancer and associated with PD-1 blockade response using spatial transcriptomics for focused regional validation (22), yet their single-cell spatial organization, multicellular neighborhood architecture, and stromal signaling mechanisms in the neoadjuvant ICB setting remained entirely uncharacterized at whole-slide single-cell resolution across a clinical ICB cohort. In melanoma neoadjuvant clinical trial correlative studies, B-cell and TLS enrichment strongly associates with pathologic response to NIVO monotherapy and IPI-NIVO combination therapy (23–25). Separately, transcriptomic profiling of NIVO-RELA trial biospecimens using a T cell-centric non-spatial approach has revealed distinct CD8+ T cell differentiation dynamics via enhanced TCR signaling despite retention of an exhaustion transcriptional profile (26,27), leaving the comparative spatial immune architecture across these regimens largely unexplored. Together, these studies establish TLS architecture and spatially organized T and B cell interactions as compelling candidate biomarkers across distinct neoadjuvant ICB regimens (28). As the field advances toward precision immunotherapy through increasingly rational and combinatorial immune-based strategies, the precise quantification of TLS structures and their cellular composition has been identified as a priority for emerging TLS-targeted therapeutics and patient stratification in clinical trials (29), yet the spatially resolved single-cell frameworks and computational pathology tools needed to define, compare, and quantify these complex immune structures reliably within the immune-dense microenvironments of ICB response remain critically lacking.

Despite these advances, fundamental questions about the single-cell spatial organization of the neoadjuvant ICB microenvironment remain unanswered: how are TCF7+ stem-like T cells spatially localized within the ICB tumor bed? What is their microenvironmental cellular and stromal context, and which chemokine axes govern multicellular immune hub assembly across distinct ICB regimens? We hypothesize that defining this spatial immune architecture and cellular interactions will identify determinants of pathologic response and resistance that differ across ICB regimens, and that these molecular discoveries can be translated into computational pathology tools to enable patient stratification and guide treatment decisions. Indeed, a recent perspective has highlighted the urgent need for well-annotated neoadjuvant ICB datasets and standardized spatial tools for TLS quantification to advance TLS-centric precision immunotherapy (29), a challenge directly addressed in this study. Beyond these two-dimensional frameworks, while early neoadjuvant ICB correlative studies have suggested TLS and germinal centers as promising histologic correlates of response (23,30–32), their three-dimensional morphologic organization (33) and the application of computational frameworks for 3D immune structure reconstruction in this context remain entirely unexplored. The emergences of subcellular resolution spatial molecular imaging (34), non-destructive 3D imaging modalities (35,36), and deep learning-based computational frameworks (37,38) now provide an unprecedented opportunity to meet these challenges, enabling multi-scale dissection of the neoadjuvant tumor microenvironment (TME) for translation into clinically actionable computational pathology tools.

To address these questions, we assembled the largest neoadjuvant immunotherapy biospecimen cohort to date for spatial multi-omics investigation, comprising 91 FFPE specimens from 87 patients with stage III melanoma, spanning three distinct ICB regimens (IPI-NIVO, NIVO-RELA, and PD-1 monotherapy) and 27 treatment-naïve controls. To systematically dissect the neoadjuvant ICB tumor ecosystem, we pursued two parallel strategies. In the first strategy, across approximately 112 million profiled cells from 105 slides, we leveraged single-cell resolution MERFISH, targeted mIF, and matched FFPE scRNA-sequencing (39) as an orthogonal validation platform to interrogate the cellular composition, immune mechanisms, and stromal networks underlying neoadjuvant ICB response and resistance. This included cellular neighborhood organization and spatially resolved receptor-ligand signaling via our novel single-cell resolution cell-cell communication algorithm for whole-slide receptor-ligand interaction quantification. We demonstrated that pathologic response is defined by the coordinated spatial assembly of a multicellular immune ecosystem centered on mature TLSs with follicular dendritic cell network and germinal centers, TCF7+ stem-like CD8 and CD4 T cells, myeloid cells, and plasma cells. We also identified CCL19+/CCL21+ fibroblasts as a previously unrecognized stromal scaffold for stem-like T cell recruitment and retention in the ICB response microenvironment. Spatially resolved receptor-ligand analyses further revealed zone-specific signaling interactions organizing the T cell, germinal center/B cell, and myeloid zones of the ICB response microenvironment, corroborating and extending the cellular neighborhood findings.

Given the central importance of TLS and germinal center structures revealed by our MERFISH analyses, we next built a discovery-to-tool pipeline for molecularly grounded TLS and germinal center quantification from spatial omics images and automated H&E-based detection. First, we leveraged MERFISH-defined molecular determinants of TLS maturation to inform low-plex mIF panel design for ICB response assessment. We then developed GCSCAN, our novel Graph-based Spatial Clustering Against Noise algorithm, to precisely delineate and quantify germinal center and TLS structures across MERFISH and mIF datasets. These molecularly validated structures served as multimodal ground truth for training PathNet-TLS, our purpose-built AI model for TLS and germinal center detection from H&E. Applying GCSCAN across the full mIF cohort, we quantified TLS and germinal center structures throughout the ICB microenvironment. Their density stratified disease-free survival, with responders lacking germinal centers at elevated risk of relapse, suggesting germinal center burden may inform long-term outcome following neoadjuvant ICB. Furthermore, we showed that IPI-NIVO induces a significantly more robust mature TLS and germinal center response than NIVO-RELA, indicating differential mechanisms of adaptive immune activation. This has direct implications for therapeutic selection and trial design. Likewise, PathNet-TLS demonstrated superior precision and interpretability compared to existing computational algorithms and pathology foundation models, addressing the urgent need for AI-based tools for standardized TLS biomarker discovery and validation in the field. Notably, PathNet-TLS orthogonally confirmed the differential mature TLS and germinal center response between IPI-NIVO and NIVO-RELA directly from H&E images, establishing a scalable and clinically actionable tool for routine H&E-based TLS and germinal center quantification. Lastly, complementing these two-dimensional analyses, open-top light-sheet imaging (35) and CODA-based 3D reconstruction of serially stacked H&E images (40) uncovered striking morphologic heterogeneity of germinal center structures in the z-dimension, including frequently interconnected TLS-GC tunnels invisible to standard 2D histopathology, with fundamental implications for current neoadjuvant response scoring practice.

In sum, this work establishes the first comprehensive single-cell resolution spatial molecular atlas of the melanoma neoadjuvant ICB ecosystem at multiple scales, introduces a discovery-to-tool paradigm linking single-cell molecular discovery to clinically deployable computational pathology tools, and lays the foundation for biomarker-driven computationally enhanced response assessment in neoadjuvant immunotherapy across multiple cancer types. As the field addresses the fundamental question of whether TLS structures represent merely correlative biomarkers or active therapeutic targets (29), the spatially resolved single-cell molecular framework, purpose-built computational tools, and neoadjuvant ICB datasets established in this study provide the foundation for resolving this question with unprecedented precision.

## RESULTS

### Study design and multimodal profiling of the neoadjuvant ICB tumor ecosystem

We assembled a cohort of 87 patients with stage III metastatic melanoma, of whom 60 received neoadjuvant ICB spanning three distinct regimens (25 IPI-NIVO, 21 NIVO-RELA, 14 PD-1 monotherapy) and 27 were treatment-naïve controls (Figure 1; Tables S1-2). All biospecimens were collected at the time of curative-intent surgery for tumor or tumor bed resection. Pathologic response assessment on H&E slides was performed by two board-certified dermatopathologists using INMC guidelines (14), based on visual estimation of residual viable tumor, necrosis, pigmented macrophages, and fibrosis within the tumor bed. Pathologic complete response (pCR) was defined as the absence of viable tumor; near-pCR as 10% or less residual viable tumor; partial response (PR) as 10–50% residual viable tumor; and non-response (NR) as greater than 50% residual viable tumor. All H&E slides underwent quality control and digital scanning followed by expert dermatopathologist annotation of tumor bed regions. A total of 91 FFPE biospecimens from these patients were then analyzed by multimodal profiling, using combinations of single-cell resolution MERFISH spatial transcriptomics (34), matched FFPE scRNA-sequencing, targeted mIF and IHC, automated H&E-based computational pathology, and non-destructive 3D imaging (35) (Figure 1A). These assays collectively profiled neoadjuvant ICB tumor microenvironments across two and three dimensions (Figure 1B).

**Figure 1.**
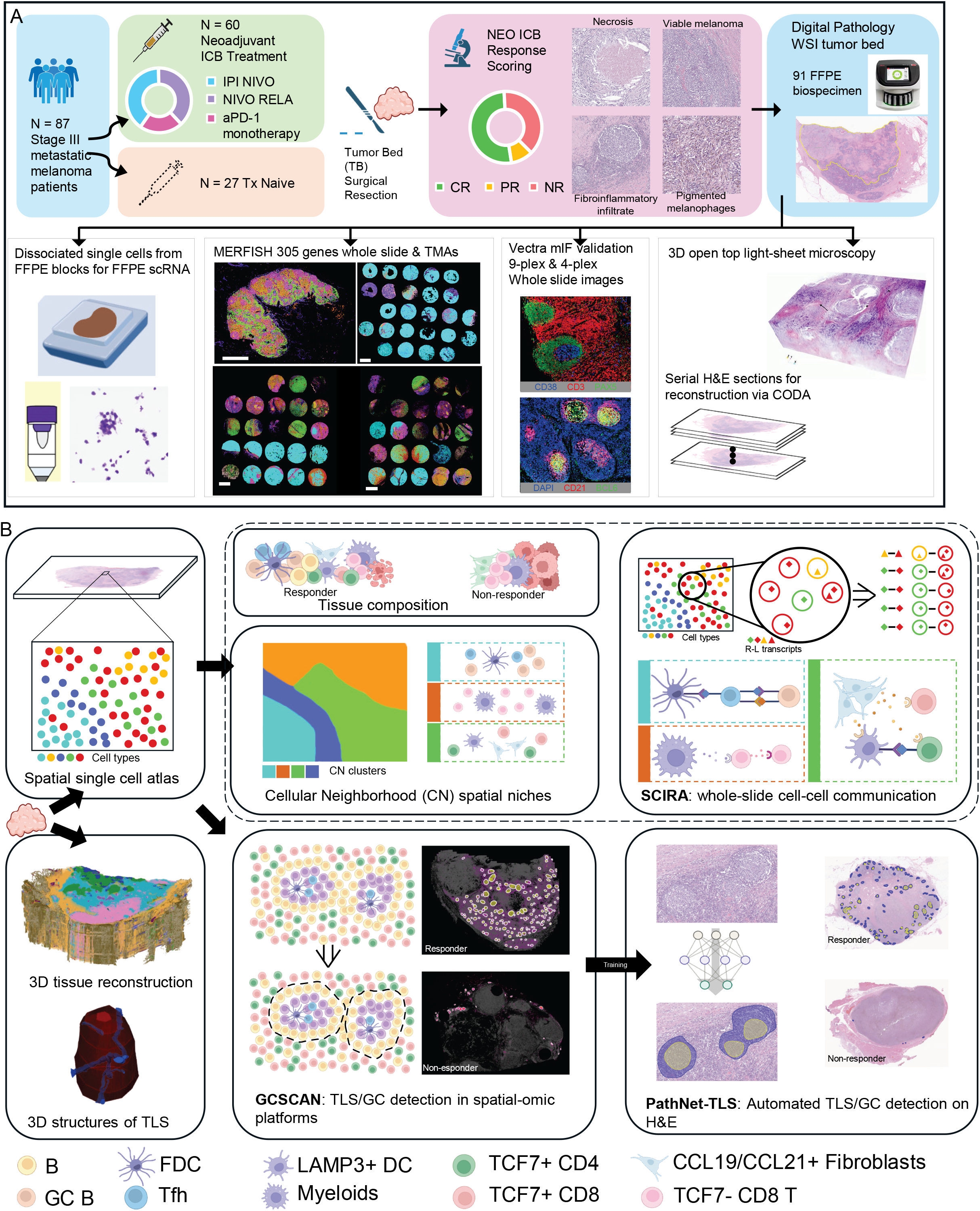
Experimental design and discovery-to-tool pipeline for comprehensive spatial profiling of the neoadjuvant ICB tumor microenvironment in 2D and 3D. (A) Study design schematic showing the patient cohort of 87 stage III melanoma patients spanning three neoadjuvant ICB regimens (IPI-NIVO, NIVO-RELA, and PD-1 monotherapy) and 27 treatment-naïve controls, INMC-based pathologic response scoring, and multi-scale spatial profiling platforms including MERFISH, Vectra mIF, FFPE scRNA-seq, 3D open-top light-sheet imaging, and CODA-based 3D computational pathology reconstruction. (B) Analytic workflow. Multimodal profiling of FFPE melanoma specimens generated a spatial single-cell atlas, through which the melanoma neoadjuvant ICB ecosystem was interrogated along three complementary analytical arms: (top) compositional tissue landscape, cellular neighborhood (CN) analysis of spatial niches, and SCIRA-based whole-slide single-cell receptor-ligand (R-L) interaction quantification; (bottom right) GCSCAN-based, molecularly grounded TLS and GC structural delineation and PathNet-TLS automated H&E-based TLS detection; and (bottom left) three-dimensional modeling of TLS and GC architecture by open-top light-sheet (OTLS) imaging and CODA-based reconstruction of serially stacked H&E sections.

### MERFISH single-cell spatial transcriptomic profiling of the neoadjuvant ICB microenvironment

To characterize the multicellular composition and cellular states of the neoadjuvant TME at single-cell resolution in situ, we performed MERFISH spatial transcriptomics using a customized 305-gene panel (Tables S3-5). We designed this panel to capture a broad spectrum of immune and non-immune tumor microenvironment cell populations, cell states including TCF7+ stem-like and immune exhaustion programs (20,41,42), critical chemokine receptor-ligand pairs, and oncogenic and metabolic pathways. We first profiled four neoadjuvant ICB-treated tumor bed whole-slide sections for two responders and two non-responders including both ICB combination regimens (NEO11: IPI-NIVO CR; NEO12: IPI-NIVO NR; NEO13: NIVO-RELA and then IPI NIVO NR; NEO14: NIVO-RELA CR). Cell type annotation was performed using a multi-step pipeline encompassing cell segmentation, quality control, high-dimensional unsupervised clustering, manual curation of marker expression, and histologic correlation (Methods), identifying 3,243,575 high-quality cells across 18 distinct cell populations: melanoma cells, B cell subsets (germinal center B cells, non-germinal center B cells, plasma cells), T cell subsets (TCF7+ stem-like CD4 and CD8 T cells, TCF7− CD4 and CD8 T cells, follicular helper T cells, and regulatory T cells), plasma cells, myeloid cells, dendritic cell subsets (classical dendritic cells (cDC), plasmacytoid dendritic cells (pDC), follicular dendritic cells (FDCs), LAMP3+ DC), endothelial cells, and 2 fibroblast subsets (Methods; Figures 2A–D; Figure S1A). We observed cell-type-specific chemokine and cytokine receptor-ligand expression patterns across these populations (Figures S1B and S1C), including CCR7 and CCL19/CCL21 in TCF7+ stem-like T cells and LAMP3+ dendritic cells; CCL5 in TCF7− CD8 T cells; CXCL13 in FDCs; and two transcriptionally distinct fibroblast subpopulations distinguished by CCL19/CCL21 expression. To orthogonally validate MERFISH cell type annotations, we performed scRNA-sequencing on the same tumor bed FFPE blocks using matched adjacent sections (39). Despite the inherently lower cell yield of scRNA due to dissociation and encapsulation constraints, comparable tumor microenvironment cell populations and cell-type-specific chemokine expression patterns were identified across both platforms (Figure 2E; Figures S2A and S2B).

**Figure 2.**
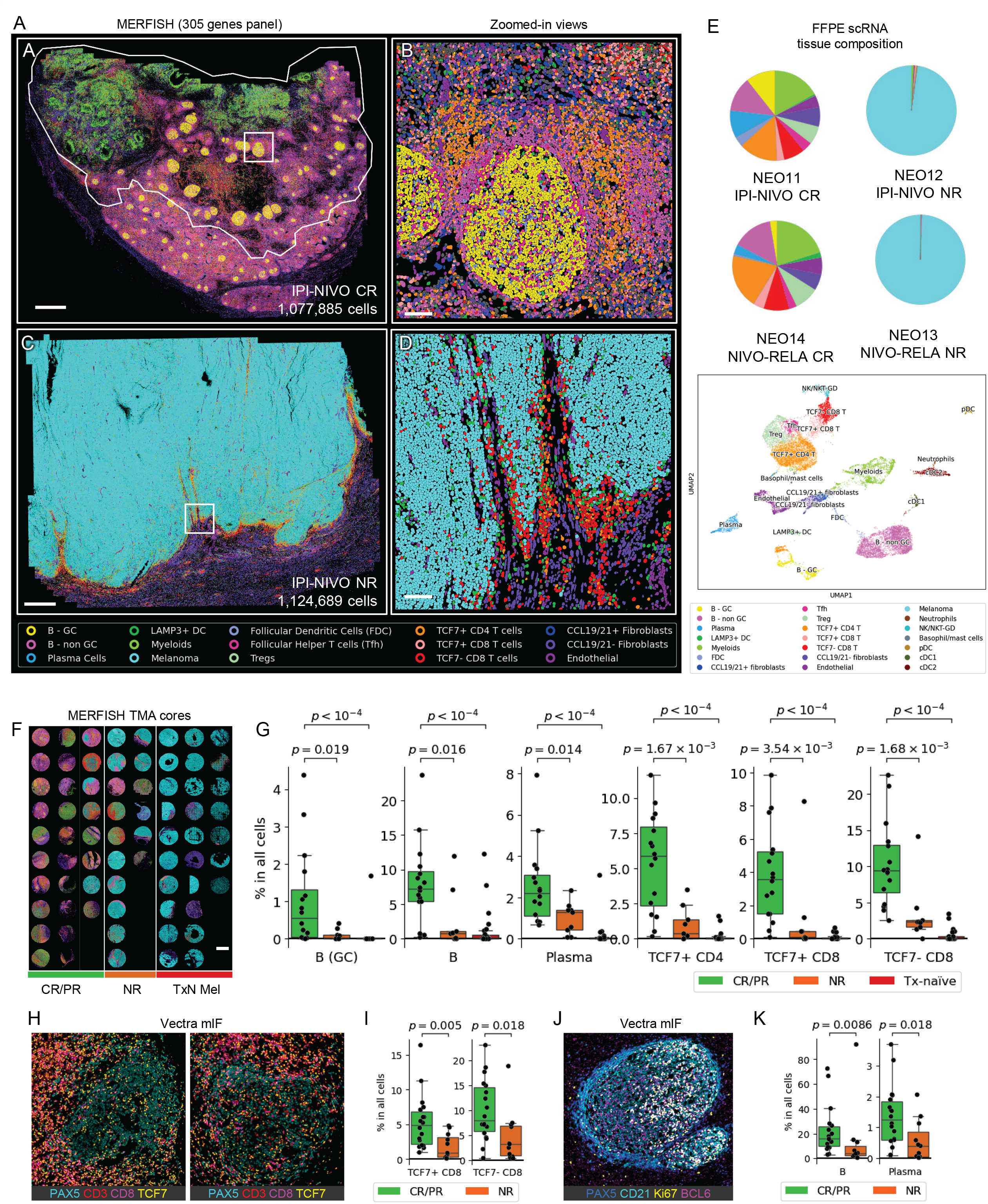
Spatial composition of the neoadjuvant ICB tumor microenvironment by MERFISH and Vectra mIF reveals differences in immune cell composition and spatial organization between responders and non-responders. (A-D) MERFISH whole-slide images (WSIs) of tumor beds from representative patients: NEO11 (IPI-NIVO CR, A-B) and NEO12 (IPI-NIVO NR, C-D). Cell type assignments are indicated by color. Color code is at bottom. (B, D) show zoomed-in views corresponding to the white rectangles in (A) and (C). White boundaries mark the annotated tumor bed. Scale bar = 1 mm (A, C) and 100 μm (B, D). (E) Cell type composition pie charts derived from FFPE scRNA-seq of the same tissue blocks for NEO11-14 (top). UMAP of 21 cell types in these samples (bottom). (F) MERFISH TMA cores sorted by response group (CR/PR, NR, treatment-naïve). Color code is the same as in A-D. Scale bar = 1 mm. Tx-naïve: treatment-naïve. (G) Boxplots comparing immune cell population proportions across response groups (n=14 CR/PR, 7 NR, and 27 treatment-naïve patients). P-values are from Mann-Whitney tests and adjusted for multiple comparisons using the Holm-Bonferroni method. (H-K) Validation of T cell and B cell immune hubs by Vectra 9-plex mIF (n=29: 18 CR/PR, 11 NR patients). (H) Representative images of T cell and B cell markers highlighting PAX5+ B cells as well as TCF7+ versus TCF7− stem-like CD8 T cells. (I) Boxplots comparing TCF7+ and TCF7− CD8 T cell proportions between response groups (Mann-Whitney test). (J) Representative image with B cell (PAX5), follicular dendritic cell (CD21) and germinal center (BCL6, Ki67) markers. (K) Boxplots comparing B and plasma cell proportions between response groups (Mann-Whitney test).

Spatial mapping of these 18 TME subpopulations across the NEO11–14 whole slide images (WSIs) revealed strikingly distinct immune landscapes between responders and non-responders. The two CR tumor beds exhibited immune-inflamed microenvironments (NEO11, Figures 2A and 2B; Figure S3; NEO14, Figures S4A and S4B), whereas the two NR tumor beds showed marked paucity of immune infiltration (NEO12, Figures 2C and 2D; NEO13, Figures S4C and S4D). The NEO11 tumor bed (Figures 2A and 2B) harbored numerous mature TLS structures containing germinal centers, Tfh cells, and FDCs. These TLSs displayed marked heterogeneity in size and morphology, many of which appeared hyper-expanded. Surrounding the germinal centers were organized zones of non-germinal center B cells and inter-germinal center T cells including TCF7+ stem-like CD4 and CD8 T cells, as well as Tregs, LAMP3+ dendritic cells, and plasma cells. Comparable cellular composition and spatial organization were observed in NEO14, though the germinal center response appeared more robust in the IPI-NIVO CR tumor bed compared to the NIVO-RELA CR in both MERFISH (Figures 2A and 2B; Figures S4A and S4B) and scRNA-seq (Figure 2E) datasets. Beyond lymphoid populations, MERFISH revealed spatially distinct CCL19/CCL21-expressing and non-expressing fibroblast subpopulations occupying distinct niches, with CCL19/CCL21-expressing fibroblasts enriched in immune-inflamed regions alongside TCF7+ stem-like T cells, and CCL19/CCL21-negative fibroblasts predominating in regions enriched for exhausted TCF7− CD8 T cells (Figure S3).

In contrast, the two NR tumor beds were dominated by melanoma cells with sparse myeloid infiltration, exhausted TCF7− CD8 T cells, CCL19/CCL21-negative fibroblasts, and endothelial networks (Figures 2C and 2D; Figures S4C and S4D). NEO12 displayed an immune exclusion phenotype characterized by a thin rim of TCF7− exhausted CD8 T cells and myeloid cells at the tumor periphery (Figures 2C and 2D), while NEO13 presented an immune desert phenotype with sparsely scattered TCF7− exhausted CD8 T cells, myeloid cells, and rare clusters of TCF7+ stem-like CD8 T cells within the tumor bulk (Figures S4C and S4D). Notably, CCL19/CCL21 expressing fibroblasts were significantly lower in non-responding tumor beds (Figures S4E and S4F), in striking contrast to their enrichment in responding microenvironments (Figures 2A–D).

To extend these findings beyond the discovery whole-slide cohort, we performed MERFISH profiling of two neoadjuvant tissue microarrays (TMAs) (n = 43 cores) and a treatment-naïve TMA (n = 27 cores), analyzing 2.36 million high-quality cells across 70 tissue cores from 48 patients (Figure 2F). Consistent with the WSI findings, the TMAs also demonstrated significant enrichment of germinal center B cells, non-germinal center B cells, plasma cells, TCF7+ stem-like CD4 and CD8 T cells, Tfh cells, TCF7− CD8 T cells, FDCs, CCL19/CCL21+ fibroblasts, myeloids, and endothelial cells in neoadjuvant ICB responders compared to non-responders and treatment-naïve controls (Figure 2G; Figure S5; Table S6). We further validated the immune-cell findings by performing targeted mIF profiling of neoadjuvant tumor bed whole-slide sections (n = 54 patients) using two low-plex antibody panels whose markers were directly informed by the MERFISH-defined cellular determinants of TLS maturation and ICB response (PAX5, CD21, Ki67, BCL6, CD38, CD3, CD8, TCF7, LAMP3; Figure S6). mIF confirmed significant enrichment of mature TLS comprising germinal center B cells and FDCs, plasma cells, and surrounding TCF7+ stem-like CD8 T cells in neoadjuvant ICB responders compared to non-responders (Figures 2H–K; Table S7).

### Cellular neighborhood analyses reveal distinct immune hubs in neoadjuvant ICB response

Having defined the critical immune and stromal cell types and states underlying neoadjuvant ICB response and resistance, we next sought to characterize the spatial molecular architecture of the tumor microenvironment through cellular neighborhood (CN) analysis at single-cell resolution (Figure 3). CN expression profiles were computed as the average gene expression among the 10 nearest neighbors of each cell and clustered into distinct CN clusters via unsupervised clustering (Figure S7A; Methods), with the spatial distribution and dominant cell populations of each CN cluster subsequently mapped and quantified. Analysis of the IPI-NIVO CR tumor microenvironment (NEO11) revealed 9 distinct CN clusters (Figures 3A–C), each defined by unique combinations of dominant cell types: germinal center light and dark zones (CN1 and CN2), a myeloid-rich zone (CN3), two T cell-rich zones (CN4 and CN8), an endothelial network zone (CN5), two fibrosis zones separately resolved into tumor bed and non-tumor bed neighborhoods based on CN expression profiles and histologic correlation with matched H&E images (CN6 and CN9), and a non-germinal center B cell zone (CN7). Importantly, CN analysis enabled both precise characterization of the cellular constituents within each neighborhood zone and reciprocal visualization of key cellular environments for each cell type (Figures 3B and 3C).

**Figure 3.**
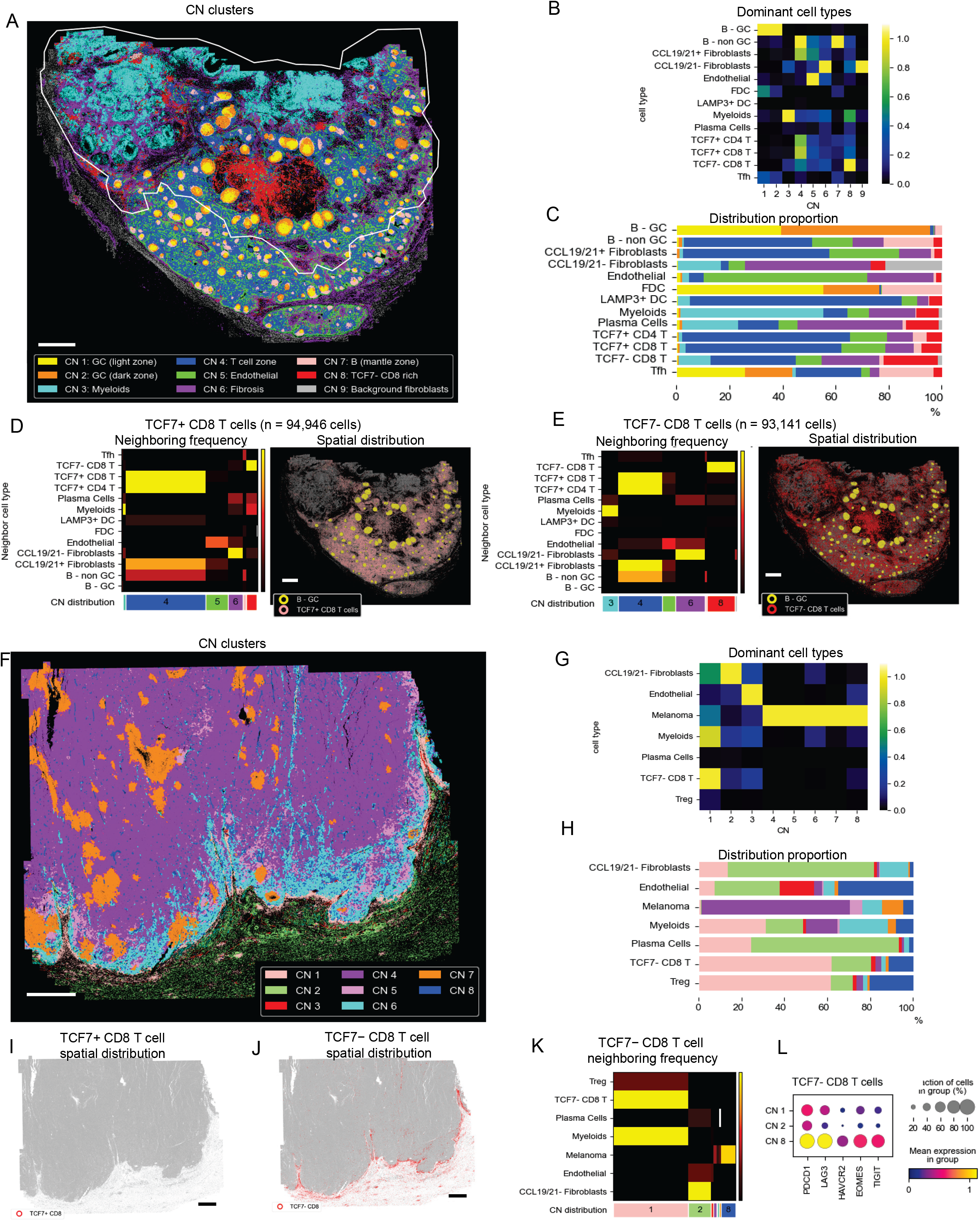
Cellular neighborhood analysis reveals distinct local immune microenvironments within the neoadjuvant ICB tumor bed. (A-E) Analysis of NEO11 (IPI-NIVO CR, WSI), comprising 1,077,885 cells. (A) CN map with white contour line highlighting the annotated tumor bed. Color code for each CN denoted at bottom. Scale bar = 1 mm. (B) Heatmap showing the dominant cell types in each CN, with color representing proportions of the cell type within each CN normalized by the maximum value per CN (column). (C) CN distribution for each cell type. (D) Characterization of TCF7+ CD8 T cells (n=94,946) and their neighborhood microenvironment. Left: heatmap showing permutation z-scores for neighboring frequency of TCF7+ CD8 T cells with other indicated cell types; columns represent CNs, with widths proportional to TCF7+ CD8 T cell distribution across CNs. Right: spatial localization of TCF7+ CD8 T cells relative to germinal centers. (E) Characterization of TCF7− CD8 T cells (n=93,141) and their neighborhood microenvironment. Left: heatmap showing permutation z-scores for neighboring frequency of TCF7− CD8 T cells with other indicated cell types; columns represent CNs, with widths proportional to TCF7− CD8 T cell distribution across CNs. Right: spatial localization of TCF7− CD8 T cells relative to germinal centers. Scale bar = 1 mm. (F-L) Analysis of NEO12 (IPI-NIVO NR, WSI), comprising 1,124,689 cells. (F) CN map with white contour line highlighting the annotated tumor bed. Color code for each CN denoted at bottom. Scale bar = 1 mm. (G) Heatmap showing the dominant cell types in each CN. (H) CN distribution for each cell type. (I-J) Spatial localization of TCF7+ CD8 T cells (I, n=0 cells) and TCF7− CD8 T cells (J, n=27,202 cells). Scale bar = 1 mm. (K) Heatmap showing permutation z-scores for neighboring frequency of TCF7− CD8 T cells with other indicated cell types; columns represent CNs, with widths proportional to TCF7− CD8 T cell distribution across CNs. (L) Dot plot showing elevated expression of T cell inhibitory checkpoint and effector genes (PDCD1, LAG3, HAVCR2, EOMES, TIGIT) in TCF7− CD8 T cells in CN 8.

While prior single-cell studies established the critical role of TCF7+ stem-like CD8 T cells in ICB response (20,22,42), their spatial localization and microenvironmental interactions have remained poorly defined. Our single-cell resolution CN analyses revealed concentration of TCF7+ CD8 T cells within CN4, the distinct T cell zone, co-enriched with TCF7+ stem-like CD4 and CD8 T cells, CCL19+ and CCL21+ fibroblasts, and non-germinal center B cells (Figures 3B and 3C). CN-specific analysis of the preferential neighbors of the TCF7+ CD8 T cells confirmed that in CN4, their preferred spatial neighbors were CCL19+/CCL21+ fibroblasts, TCF7+ stem-like CD4 and CD8 T cells, and non-germinal center B cells (Figure 3D). In contrast, TCF7− exhausted CD8 T cells were distributed more broadly across CN4, CN6, and CN8 (Figures 3B and 3C). These CNs exhibited distinct co-enriched cell populations: CN4 with TCF7+ stem-like CD4 and CD8 T cells, CCL19+/CCL21+ fibroblasts, and non-germinal center B cells; CN6 with CCL19/CCL21− fibroblasts; and CN8 with other exhausted TCF7− CD8 T cells, myeloid cells, and CCL19/CCL21− fibroblasts (Figure 3B). Analysis of TCF7− CD8 T cell CN-specific preferential neighbors revealed neighboring patterns consistent with the co-enrichment findings (Figure 3E). Although TCF7− CD8 T cells and myeloids show noticeable co-enrichment in CN8 (Figure 3B), their neighboring tendency is strongest in CN3 (Figure 3E). Additional notable CNs include the germinal center light zone (CN1) enriched for germinal center B cells, FDCs, and Tfh cells; the endothelial-rich CN (CN5) co-enriched with non-germinal center B cells and CCL19/CCL21+ fibroblasts; and the myeloid CN (CN3) enriched for CCL19/CCL21−fibroblasts (Figures 3B and 3C). Analogous CN profiling of the NIVO-RELA CR tumor microenvironment identified 9 comparable CN clusters with similar dominant cell types and preferential neighboring frequencies (NEO14; Figures S7B–F).

Spatial CN profiling of the NR tumor microenvironments revealed a distinct landscape. NEO12 (IPI-NIVO NR) yielded 8 CN clusters, five of which were occupied predominantly by melanoma cells (CN4–8; Figures 3F–H). Despite the overall paucity of immune infiltration in this immune-excluded microenvironment, CN1 showed co-enrichment of TCF7− exhausted CD8 T cells, myeloid cells, CCL19/CCL21−fibroblasts, and a subset of melanoma cells. The remaining CNs were characterized by enrichment of CCL19/CCL21−fibroblasts (CN2) and endothelial cells (CN3). While TCF7+ stem-like CD8 T-cells are sparsely distributed within the tumor bed, a thin rim of TCF7− CD8 T cells surround the tumor bulk (Figures 3I and 3J). In CN1, these TCF7− CD8 T cells’ preferred neighbors were myeloid cells, Tregs, and other TCF7− CD8 T cells. However, in CN8, the most common neighbors of TCF7− CD8 T cells were melanoma cells (Figures 3G, 3H, and 3K). Notably, TCF7− CD8 T cells in CN8 appeared more exhausted than those in CN1 (Figure 3L). In the immune desert NEO13 tumor microenvironment (NIVO-RELA NR; Figures S7G–I), spatial CN profiling revealed only 4 CN clusters, all dominated by melanoma cells. Similar to NEO12, the TCF7−exhausted CD8 T cells are co-enriched with CCL19/CCL21− fibroblasts, endothelial cells, myeloid cells, and plasma cells in CN4. Strikingly, while TCF7+ CD8 T cells were scattered within these NR tumor microenvironments, they did not form meaningful CN clusters in either NEO12 (Figures 3G and 3I) or NEO13 (Figures S4C and S4D; Figure S7I). This was in marked contrast to their organized spatial enrichment within the ICB responder tumor microenvironments (NEO11, NEO14). Together, these CN analyses reveal that organized spatial assembly of germinal center B-cells and TCF7+ stem-like T cells with CCL19/CCL21+ fibroblasts and non-germinal center B cells is a defining feature of the neoadjuvant ICB response microenvironment absent in non-responding tumor beds.

### Cell-cell communication in neoadjuvant ICB response

In the spatially organized TME, cell identity is shaped not only by cell-autonomous gene expression but also by non-autonomous signals from neighboring cells. Building on the distinct cellular neighborhoods identified above, we next sought to elucidate the molecular interactions between critical cell types underlying neoadjuvant ICB response. Application of existing receptor-ligand interaction methods to our whole-slide MERFISH datasets revealed fundamental constraints. COMMOT (43), a widely-used spatially-aware single-cell resolution receptor-ligand computational method based on collective optimal transport, exhibited prohibitive scaling of computation time and memory requirements with increasing cell numbers, failing entirely at the scale of the millions of cells in WSIs (Figure S8). Its analysis was therefore constrained to predefined immune hub regions rather than unbiased whole-slide analyses, as previously employed in lung cancer spatial transcriptomics (22), precluding discovery of novel cell-cell interactions outside the predefined zones. CellChat v2 (44,45) offers improved scalability, but it operates at the cell group level, precluding true single-cell pairwise receptor-ligand quantification. To overcome these limitations, we developed a novel computational method, Spatial Cellular Interaction and Receptor Activation (SCIRA), to quantify statistically significant receptor-ligand (R-L) signaling between pairs of cells at the whole-slide scale (Figure 4A). SCIRA models the degree of receptor activation for each cell based on the cell’s receptor expression and the total ligand available from neighboring cells within a defined local radius (15 µm), following simplified mass action kinetics (Figure 4A; Methods). This design enables efficient and unbiased single-cell level R-L interaction quantification across millions of cells in whole-slide images and TMAs. When benchmarked against COMMOT, SCIRA demonstrated significantly faster computation speed and lower memory requirements than COMMOT (Figures S8A and S8B) while generating comparable R-L interactions in the neoadjuvant TMA (Figure S8C). Importantly, it completed unbiased whole-slide level R-L interaction quantification within 500 seconds in datasets where COMMOT failed due to memory limitations.

**Figure 4.**
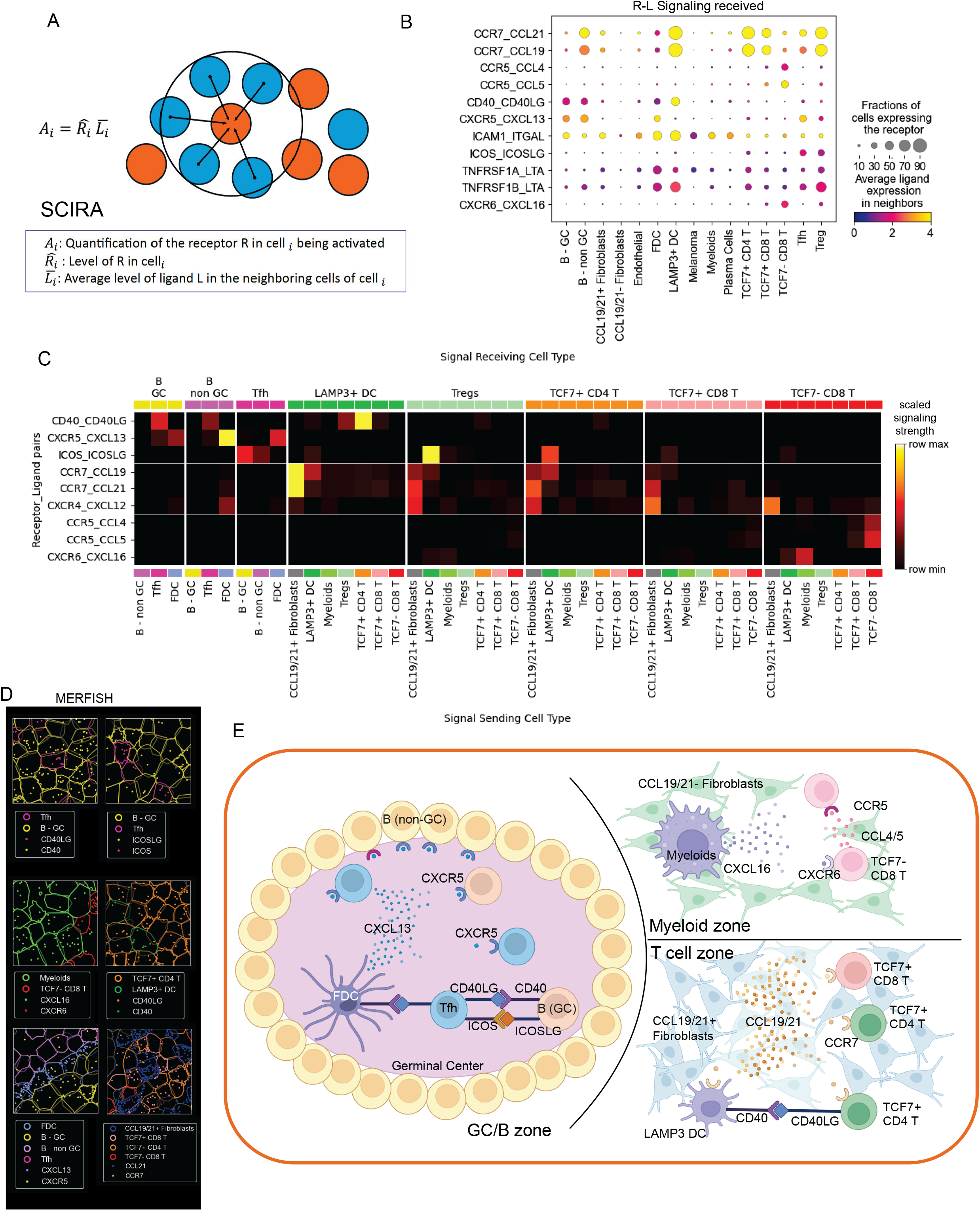
Spatially resolved receptor-ligand signaling interactions in neoadjuvant ICB response revealed by SCIRA. (A) Schematic illustrating the SCIRA algorithm, which quantifies cell-cell receptor-ligand signaling based on receptor expression levels and ligand availability in neighboring cells within a defined radius (15 µm). (B) Dot plot showing top-ranked receptor-ligand signaling interactions received by each cell type (x-axis). Dot size indicates fraction of cells expressing the receptor; color intensity indicates cell-type-average total ligand expression in neighboring cells. Shown data are from MERFISH whole-slide and TMA datasets. (C) Heatmap showing scaled signaling strength for receptor-ligand pairs (rows) between signal-sending and signal-receiving cell type pairs (columns), highlighting preferential interactions within the T cell, germinal center/B cell, and myeloid zones. Shown data are from MERFISH whole-slide and TMA datasets. (D) Zoomed-in MERFISH view showing subcellular RNA detection and cell boundaries colored by cell type, with representative insets illustrating spatial co-localization of signal-sending and signal-receiving cell pairs for representative receptor-ligand interactions. (E) Schematic summarizing key intercellular receptor-ligand signaling interactions and interacting cell types in neoadjuvant ICB response, including cell-cell communication within the T cell zone, germinal center/B cell zone, and myeloid zone.

Using SCIRA, we screened 45 receptor-ligand pairs across 15 cell types in our MERFISH WSI and TMA datasets, identifying the most prominent R-L interactions for each signal-receiving cell type in neoadjuvant ICB response (Figure 4B). For each signal-receiving cell, we then interrogated the signal-sending cell or cells for each R-L pair of interest (Figure 4C; Methods; Table S8). This analysis revealed preferential R-L interactions within the distinct CN zones identified above, specifically the T cell zone, germinal center/B cell zone, and myeloid zone (Figures 4D and 4E). In the T cell zone, SCIRA revealed preferential CCR7-CCL19/CCL21 interactions between TCF7+ stem-like CD8 T cells and CCL19/CCL21+ fibroblasts, corroborating the spatial co-enrichment identified by CN analysis. It also revealed CD40-CD40L interactions between LAMP3+ dendritic cells and TCF7+ stem-like CD4 T cells. In the germinal center/B cell zone, germinal center B cells and follicular helper T cells showed preferential interactions via CD40-CD40L and ICOS-ICOSL, and germinal center B cells and follicular dendritic cells interacted preferentially via CXCR5-CXCL13. In the myeloid zone, preferential interactions between myeloid cells and TCF7− CD8 T cells via CXCR6-CXCL16 were identified. Homotypic interactions among TCF7− CD8 T cells via CCR5-CCL4/CCL5 were also detected. The signal-sending and signal-receiving cell pairs identified within the T cell, germinal center/B cell, and myeloid zones are consistent with the dominant cell types and neighboring frequencies observed in CN analyses, providing independent receptor-ligand level validation of the spatially organized immune hubs characterizing neoadjuvant ICB response. Together, SCIRA analyses reveal a zonally structured intercellular communication network in which distinct chemokine and co-stimulatory signaling circuits operate within the T cell, germinal center/B cell, and myeloid zones of the ICB response microenvironment (Figure 4E).

### GCSCAN detects differential TLS responses across combination neoadjuvant ICB therapies

Having characterized the intercellular communication networks organizing the neoadjuvant ICB immune response, we next sought to quantify TLS and germinal center structures as discrete spatial entities across the full mIF cohort. While our MERFISH, scRNA-seq, and mIF analyses have collectively established TLS and GC structures as central features of the neoadjuvant ICB response microenvironment, these approaches could not delineate or quantify GC and TLS structures as organized molecular immune hubs, a prerequisite for systematic and unbiased quantification across patients and response groups. Clinically, current neoadjuvant ICB response assessment relies on visual estimation of residual viable tumor and fibroinflammatory stroma on H&E slides (14,16), a method that is inherently imprecise and fails to leverage TLS and GC structures as quantifiable immune biomarkers. Despite their striking association with ICB response established above, TLS and GC structures have never been incorporated into neoadjuvant response scoring, partly due to the absence of computational algorithms capable of reliably delineating individual GC and TLS boundaries within the immune-dense and morphologically complex microenvironments of ICB response.

To address this, we first benchmarked existing unsupervised clustering algorithms for GC and TLS segmentation in our MERFISH and mIF datasets, including Louvain and Leiden clustering (46) and the density-based algorithm Hierarchical Density-Based Spatial Clustering of Applications with Noise (HDBSCAN) (47). While HDBSCAN demonstrated superior performance compared to Louvain and Leiden clustering, it failed to accurately partition individual GC and TLS structures in the immune-dense ICB tumor microenvironment, a prerequisite for precise quantification (Figures S9A–C). To overcome this limitation, we developed GCSCAN (Figure S9D; Methods), utilizing a graph-based spatial clustering algorithm incorporating biological spatial cellular organization and graph computations to delineate GC, TLS, and B cell follicle structures from spatial omics imaging data with respect to background noise. GCSCAN demonstrated superior partitioning and identification of individual GC and TLS structures compared to HDBSCAN (Figure S9C). Importantly, GCSCAN is generalizable across spatial omics platforms for the detection of clustered immune structures. By deriving structural boundaries from MERFISH-defined single-cell molecular identities and mIF protein expression signals, rather than from morphology-based visual annotation or unsupervised clustering, GCSCAN eliminates the annotation noise and biological ambiguity inherent to existing approaches. This replaces subjective structural assignment with objective, molecularly validated boundaries at single-cell resolution.

We next applied GCSCAN to quantify GC and TLS structures across the neoadjuvant MERFISH and mIF datasets (n = 58 patients; examples shown in Figure 5A; Table S9). GCSCAN quantification revealed that the number of mature TLS and B-cell follicles and the upper quantile of GC area per mature TLS were markedly elevated in neoadjuvant ICB responders compared to non-responders (Figure 5B). Importantly, we found that IPI-NIVO responders were associated with a significantly greater number of mature TLS and GCs compared to NIVO-RELA responders (Figure 5C). This structural difference suggests that these combination ICB regimens may differentially engage germinal center-associated adaptive immunity, a distinction that has not previously been resolved at this scale. This may inform future therapeutic selection and neoadjuvant trial design. Importantly, GCSCAN-quantified GC and TLS abundance each stratified disease-free survival in neoadjuvant ICB-treated patients (n = 58 with MERFISH or mIF WSI; Figures 5D and 5E; log-rank p = 0.0204 for TLS and p = 0.0019 for GC). We followed these patients over time. Three of four responders who subsequently progressed were below the cohort median for GC density, with minimal or absent GC, at the time of curative-intent surgery (Figure 5F). This suggests that quantification of GC/mature TLS at the time of neoadjuvant curative-intent surgery may carry important prognostic information for long-term outcomes, and that the absence of organized, mature TLS/GC immune structures can herald the risk of relapse or progression.

**Figure 5.**
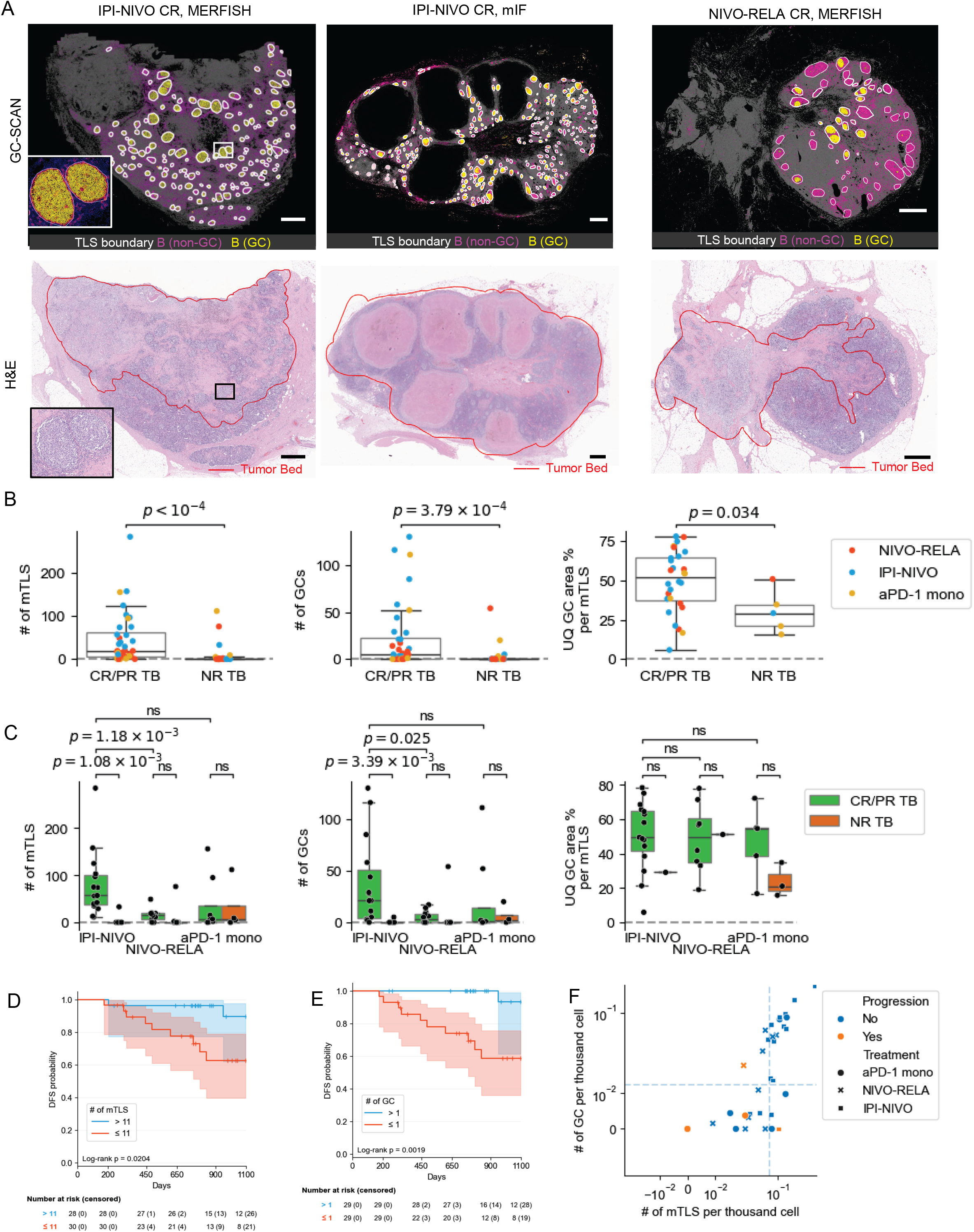
GCSCAN precisely delineates and quantifies TLS and germinal center structures across spatial omics and mIF imaging datasets, revealing distinct responses across treatment regimens. (A) Representative whole-slide images demonstrating GCSCAN-based structural delineation of mature TLS (mTLS; B cell follicle with CD21+ follicular dendritic cell network) and germinal centers (GC) in IPI-NIVO CR (left and middle columns) and NIVO-RELA CR (right column) tumor beds. (Top) MERFISH or mIF cell type map (non-GC B cells, magenta; GC B cells, yellow) with white GCSCAN-delineated mTLS boundaries overlaid on the image. Inset shows zoomed-in GC and mTLS structures; (bottom) corresponding H&E whole-slide images with tumor bed annotated in red. Scale bar = 1 mm. (B-C) Boxplots comparing GCSCAN-quantified mTLS and GC metrics across the full mIF cohort. (Left) total number of mTLS per TB; (middle) number of GCs per TB; (right) upper quartile (UQ) GC area percentage per mTLS. Each dot represents one tumor bed. (B) Comparison between responder (CR/PR) and non-responder (NR) tumor beds (n=36 CR/PR and n=21 NR), colored by treatment regimen. (C) Metrics stratified by treatment regimen and response group (green: CR/PR; orange: NR). For the area ratio comparison in both panels, only tumor beds containing GCs are considered (n=28 CR/PR and n=5 NR). Mann-Whitney test; p values adjusted for multiple comparisons using the Holm-Bonferroni method. ns, p > 0.05. (D-E) Kaplan-Meier curves showing disease-free survival (DFS) for patients with high (blue) versus low (red) mTLS (D) or GC (E) counts, dichotomized at the cohort median. Shaded bands represent 95% confidence intervals. Curves are truncated at 3 years. Numbers at risk, with cumulative censored patients in parentheses, are shown below the plot. P-values are from log-rank test. (F) Scatter plot showing distribution of mTLS (x) and GC (y) density for patients. Marker style represents treatment regimen. Orange color represents disease progression. Dashed lines mark cohort median values.

### PathNet-TLS: an end-to-end AI algorithm for automated TLS and germinal center detection from H&E

Recent advances in computational pathology have generated immense interest in identifying predictive and prognostic biomarkers of ICB response from routine H&E whole-slide images. The multi-resolution deep learning model HookNet-TLS has recently been highlighted for its ability to automatically detect TLS and germinal centers across multiple cancer types (48,49). Histopathology foundation models pretrained on large-scale WSI datasets, including UNI, GigaPath, and C-TransPath (38,50,51), have demonstrated broad capability in histological feature extraction and cancer prognostication. Nevertheless, the performance of these models in immunotherapy cohorts remains largely unexplored, and limitations in foundation model interpretability continue to pose challenges for real-world clinical implementation. Given that H&E slides acquired during routine clinical care provide the most cost-effective visualization of tissue architecture and cellular morphology at high resolution, we first evaluated the utility of HookNet-TLS and current histopathology foundation models on our neoadjuvant ICB cohort.

To establish a molecularly validated multimodal ground truth for TLS and germinal center structures, we leveraged matched H&E, mIF, and IHC whole-slide images from our neoadjuvant ICB cohort (Figure S10). TLS and germinal center structures are notoriously difficult to define by manual annotation without immune biomarkers, and existing models trained on such annotations inherit their ambiguity (49,52). To overcome this, we selected H&E slides enriched for germinal centers and performed de-staining and re-staining with Ki67/CD21 dual-plex IHC on the same tissue sections, enabling germinal center identification with direct spatial correspondence to the H&E morphology on the same tissue section (1,170 GCs, 18 slides). Mature TLSs harboring follicular dendritic cells (TLS-FDC) were characterized by co-registering matched mIF data (CD3/PAX5/CD21/Ki67) to the H&E images (993 TLS, 13 slides). GC and TLS structures in the mIF and IHC datasets were then delineated using GCSCAN, providing molecularly validated structural boundaries that replace subjective visual annotation with objective, single-cell resolution ground truth. Critically, each step in this multimodal annotation chain, from MERFISH-informed mIF panel design through GCSCAN-based structural delineation to H&E implementation, removes a layer of annotation noise and biological ambiguity, yielding a ground truth of unprecedented precision for computational pathology model training.

When benchmarked against this multimodal ground truth, HookNet-TLS failed to adequately delineate germinal centers and mature TLSs with FDCs in our neoadjuvant ICB cohort (Figure 6A). Large regions of the tumor bed were incorrectly labeled as confluent TLS without discrete follicular boundaries, and individual germinal center structures were often not accurately resolved. This suboptimal performance likely reflects two intrinsic limitations of the HookNet-TLS training cohort: 1) underrepresentation of slides with dense lymphoid stroma, mature TLS, and germinal center structures in the ICB setting, and 2) reliance on manual annotations that are often inaccurate particularly in immune-enriched microenvironments. These limitations constrain generalization to the immunologically complex neoadjuvant ICB tumor microenvironment. In contrast, patch-level TLS classifiers trained on UNI-derived patch embeddings with GCSCAN-derived TLS ground truth (Methods) produced predictions that more closely approximated TLS regions (Figure 6B). However, even the best-performing classifier still frequently failed to distinguish discrete TLS structures from surrounding non-TLS immune stroma. This indicates that even with spatial context and supervision from molecularly defined labels, patch-level foundation model embeddings may lack the discriminative features to separate TLSs from morphologically similar but functionally distinct immune aggregates.

**Figure 6.**
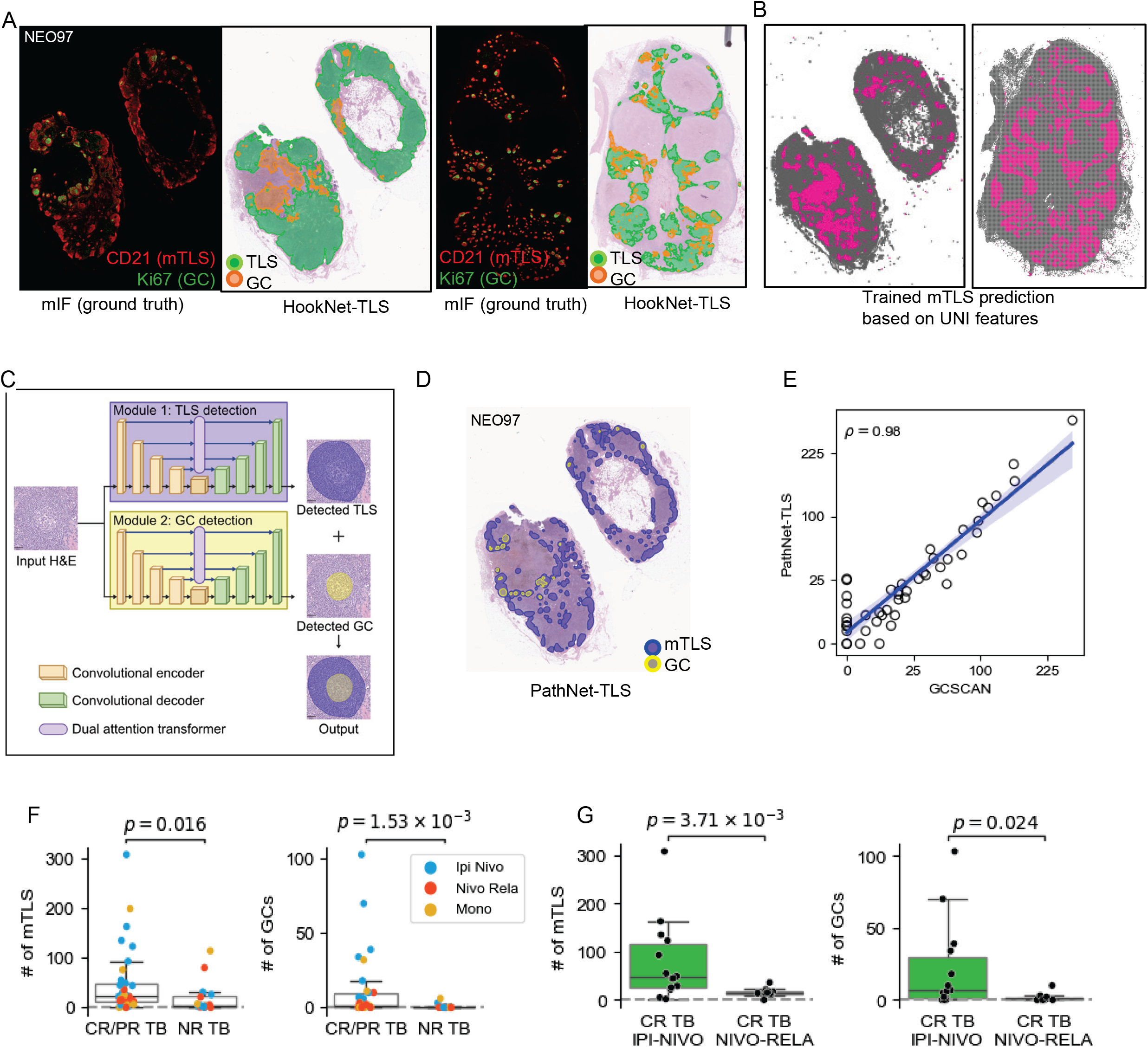
PathNet-TLS enables precise automated detection of TLS and germinal centers from routine H&E whole-slide images in real-world immunotherapy settings. (A) Side-by-side comparison of ground truth mIF and HookNet-TLS-predicted TLS and germinal center boundaries overlaid on H&E images. In mIF, ground truth shown with CD21 (red) and Ki67 (green). In HookNet-TLS overlays, green shows predicted TLS and orange GCs. (B) Visualization of TLS classification by a trained patch-level TLS classifier using UNI patch encoding with contextual feature aggregation. Magenta: patches predicted positive for TLS; Gray: patches predicted negative for TLS. (C) PathNet-TLS consists of a TLS module and a GC module. (D) Overlay of PathNet-TLS-predicted TLS and GC boundaries on a representative H&E whole-slide image. (E) Correlation plot between the number of TLS per sample predicted by PathNet-TLS and that quantified from mIF using GCSCAN, across H&Es. Axes are plotted on a square root scale. Each dot represents a tumor bed. Blue line shows linear regression and 95% interval in the square root space. Pearson correlation coefficient in original scale = 0.98; in square root scale = 0.92. (F) Boxplot showing that responders have significantly more mTLS (left, p = 0.016) and germinal centers (right, p = 0.00153) than non-responders based on PathNet-TLS quantification Mann-Whitney test. n=33 CR/PR and 16 NR tumor beds (TB). (G) Boxplot showing that IPI-NIVO responders have significantly more mTLS (left, p = 0.00371) and germinal centers (right, p = 0.024) than NIVO-RELA responders in the tumor beds, based on PathNet-TLS quantification. n=14 IPI-NIVO responders and 11 NIVO-RELA responders.

To overcome these limitations, we developed PathNet-TLS, a novel end-to-end computational pathology model trained on our neoadjuvant ICB H&E and multimodal imaging data. PathNet-TLS implements UDTransNet, a transformer-based U-Net architecture incorporating dual attention transformer mechanisms for semantic segmentation(53), which has been shown to outperform both CNN-based and transformer-based segmentation models. The model comprises two task-specific modules (Figure 6C): a TLS with follicular dendritic cell network module trained on annotations derived from co-registered mIF data, and a germinal center module trained on curated H&E images with matched CD21/Ki67 dual-plex IHC from the same tissue sections (Methods).

When benchmarked against the ground truth, PathNet-TLS reliably identified germinal centers and TLS with follicular dendritic cell networks, including accurate boundary delineation, in our neoadjuvant ICB samples (Figure 6D). Mature TLS counts predicted by PathNet-TLS. were highly correlated to MERFISH and mIF-derived counts with GCSCAN (Figure 6E; Table S10). PathNet-TLS also outperformed both HookNet-TLS and UNI in identifying germinal centers and mature TLS within immune-enriched microenvironments (Figures 6A, 6B, and 6D), despite both PathNet and UNI-based classifiers sharing the same ground truth training set. This implies our model, trained end-to-end on the ground truth, learned task-specific features for delineating fine-grained TLS and germinal center boundaries in real-world ICB-treated tumor microenvironments, even where current general-purpose patch embeddings fail. Importantly, PathNet-TLS detected significantly higher mature TLS and germinal center counts in neoadjuvant ICB responders compared to non-responders (Figure 6F; p = 0.016 for mTLS and p = 0.00153 for GC). It also demonstrated that IPI-NIVO induced a more exuberant mature TLS and germinal center response compared to NIVO-RELA and PD-1 monotherapy (Figure 6G; p = 0.00371 for TLS and p = 0.024 for GC), consistent with mIF-based quantification by GCSCAN in this cohort. Collectively, PathNet-TLS completes the discovery-to-tool pipeline established in this study: from MERFISH-defined molecular determinants of TLS maturation through GCSCAN-based structural quantification to automated H&E-based detection, enabling robust and scalable quantification of TLS and germinal center responses directly from routine H&E slides in real-world immunotherapy settings. This end-to-end framework translates single-cell molecular discoveries into a clinically actionable computational pathology tool, addressing a critical gap in neoadjuvant ICB response assessment.

### 3D atlas of the neoadjuvant ICB tumor ecosystem by open-top light-sheet imaging and CODA

Recent multiplexed 3D immune profiling of colorectal cancer demonstrated that TLS structures can be interconnected in three dimensions, with spatial features suggestive of graded molecular properties (33). To investigate whether similar architectural complexity characterizes the organization of the neoadjuvant ICB tumor microenvironment in 3D, we performed open-top light-sheet (OTLS) imaging (35,54) of IPI-NIVO tumor beds (1 CR and 1 NR) after whole tissue block bleaching, clearing, and computational staining using TO-PRO3 and eosin (Figures 7A–D). Three-dimensional optical sections of the CR tumor bed (1,800–2,000 Z-stacks; 500–1,650 μm depth) revealed striking morphologic heterogeneity across Z-planes, including numerous hyper-expanded and interconnected germinal centers (Figures 7A–D; Supplemental Videos S1– 3) that appeared spatially distinct when observed in 2D (Figure 7B). To enable correlation of 3D features with molecular assays, we benchmarked histologic processing of post-OTLS imaged samples for reverse clearing and achieved clinical-grade H&E and IHC staining (CD3, CD8, CD20, CD21, Ki67, CD68) for parallel evaluation of B cell, T cell, and follicular dendritic cell structures in serial sections (Figure 7D).

**Figure 7.**
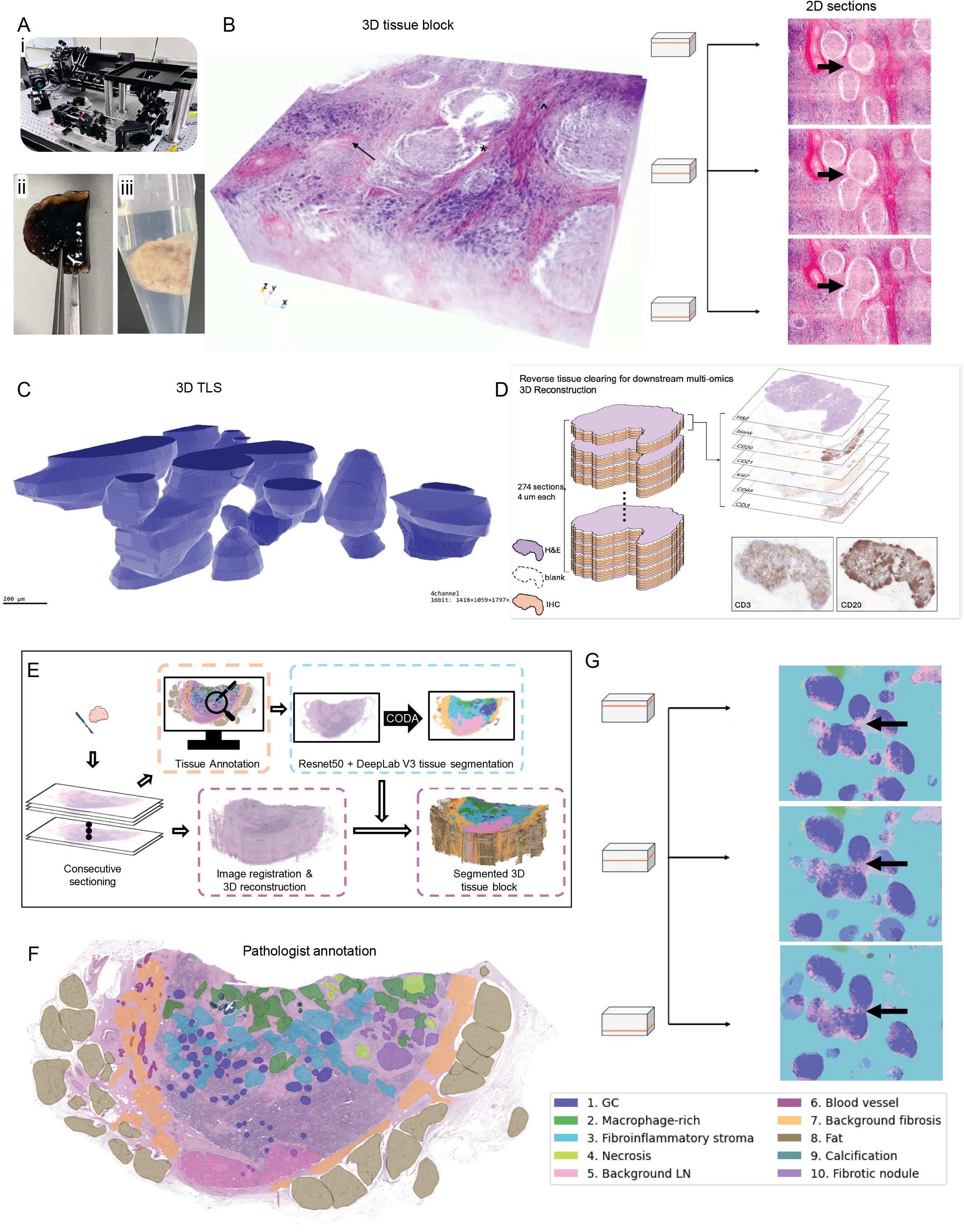
3D imaging and reconstruction of the neoadjuvant ICB tumor ecosystem reveals morphologic heterogeneity in the Z-dimension and interconnected germinal center-TLS tunnels in pathologic responders. (A) Open-top light-sheet (OTLS) microscopy. (i) OTLS microscope. (ii and iii) Representative neoadjuvant ICB surgical resection FFPE tissue block before (ii) and after (iii) bleaching and clearing. (B) OTLS 3D tissue volume rendering of an IPI-NIVO CR tumor bed (NEO11). 3D rendered tissue volume (left) and representative optical sections in Z-planes, showing that germinal centers that appeared spatially distinct in 2D are interconnected in 3D. (C) 3D rendering showing germinal center and TLS structures forming interconnected tunnels in 3D. Scale bar = 200 μm. (D) After OTLS imaging, tissues were reverse-cleared and processed for re-embedding, serial sectioning, and H&E and IHC staining. Serial H&E and CD3, CD20, and Ki67 immunostains confirm correspondence between germinal center architectures identified by OTLS and serial tissue sections. (E) Workflow for CODA-based 3D reconstruction of the neoadjuvant ICB tumor microenvironment from serial H&E image stacks, encompassing manual tissue annotation, DeepLabV3-based tissue segmentation, image registration, and 3D reconstruction of the segmented tissue block. (F) Representative H&E image showing expert pathologist annotations across 10 histologic tissue categories used for training the semantic tissue segmentation model. (G) Z-sections from the segmented reconstructed 3D tissue block, with arrows indicating interconnected germinal centers across Z-planes.

In parallel to light-sheet imaging, we implemented CODA (40), a deep learning model for 3D reconstruction of tissue microanatomy from serial H&E images, and applied it to the neoadjuvant ICB CR tumor bed (Figure 7E). We manually annotated a set of serial H&E sections (Figure 7F; NEO11, IPI-NIVO CR) and trained a deep learning semantic segmentation model for multi-label tissue reconstruction in this tumor microenvironment. CODA independently corroborated the OTLS findings, demonstrating morphologic heterogeneity across Z-planes and interconnected germinal centers in three dimensions (Figure 7G). Together, these two complementary 3D imaging and computational pathology approaches generated, for the first time, a three-dimensional histologic atlas of the melanoma neoadjuvant ICB tumor ecosystem, revealing morphologic features of immune hub connectivity suggestive of long-range molecular immune signaling invisible to standard 2D histopathology. The marked morphologic heterogeneity observed across Z-planes has direct implications for the current clinical practice of neoadjuvant response scoring based on single 2D tissue sections, underscoring the need for 3D-informed approaches to fully characterize the immune architecture of ICB response.

## DISCUSSION

Neoadjuvant immunotherapy represents one of the most important breakthroughs in cancer therapy in the past decade and a leading frontier of clinical and translational research. Central to this transformation is the adoption of pathologic response as a surrogate endpoint for trial approval across multiple cancer types, creating an unprecedented opportunity to interrogate the immune mechanisms of response from previously rare on-treatment tumor bed specimens now collected as part of routine neoadjuvant care. Yet while early correlative studies established B cell and TLS enrichment as hallmarks of ICB response, the spatial molecular architecture of the neoadjuvant ICB tumor microenvironment has remained largely undefined, as the single-cell spatial technologies needed to map it have only recently emerged. The biomarker and computational tools needed to quantify these complex immune structures and translate molecular discoveries into clinically actionable metrics have likewise been lacking.

In this study, we assembled the largest neoadjuvant immunotherapy biospecimen cohort to date and leveraged state-of-the-art single-cell spatial multi-omics, FFPE scRNA-sequencing, and 3D imaging alongside purpose-built computational tools to address these gaps, generating a comprehensive molecular and computational atlas of the melanoma neoadjuvant ICB tumor ecosystem across multiple scales. This work revealed novel hallmarks of the neoadjuvant ICB tumor microenvironment, including previously unrecognized immune hubs and stromal niches, spatially resolved intercellular signaling, and regimen-specific differences in germinal center and mature TLS induction. Importantly, we established a discovery-to-tool paradigm translating these findings into clinically deployable TLS and germinal center quantification tools that complement current histopathologic response assessment, addressing a critical unmet need in neoadjuvant ICB response evaluation. These neoadjuvant ICB biospecimens, collected as part of routine care, offer an irreplaceable resource for biomarker discovery and validation.

Tertiary lymphoid structures have been recognized as organized intratumoral immune aggregates associated with favorable prognosis and antitumor immunity across multiple cancer types (28,29), with early work demonstrating that a 12-chemokine gene expression signature encompassing CCL19 and CCL21 could identify these structures and predict patient survival across multiple solid tumor types (55,56). These early observations laid the conceptual foundation that TLS and germinal centers are key immune hubs within the tumor microenvironment, yet their spatial molecular organization at single-cell resolution, intercellular signaling, and regimen-specific induction in the neoadjuvant ICB setting have remained poorly understood. Using single-cell resolution spatial omics, we defined the full multicellular composition and cellular neighborhood organization distinguishing responders from non-responders, including the enrichment of TCF7+ stem-like T cells and CCL19/CCL21-expressing fibroblasts within organized immune hubs, and highlighting mature TLS and germinal centers as dominant features of the neoadjuvant ICB response microenvironment not captured under the current INMC guidelines. This underscores the urgent need for standardized spatial quantification tools to incorporate these immune structures into neoadjuvant ICB response assessment and advance TLS-based biomarker development, patient stratification in clinical trials, and TLS-directed precision immunotherapy.

Despite the recognition that TLS play critical roles in antitumor immunity and response to ICB, they are notoriously difficult to delineate in histopathology and spatial omics images as intact immune structures. While individual cell types can be enumerated with the assistance of biomarkers, systematic quantification of TLS as organized structural entities across patients and treatment cohorts remains virtually impossible without purpose-built computational tools. The recent pan-cancer TLS atlas (48), for example, relied on pixel-level TLS signature score thresholding derived from computationally enhanced spot-level Visium spatial transcriptomic data, without cell segmentation or spatially and individually resolved cell types, precluding precise delineation of individual TLS structures, particularly in confluent high-score regions characteristic of immune-enriched tumor microenvironments.

To our knowledge, this study also represents the first spatial omics characterization of the neoadjuvant NIVO-RELA treated tumor microenvironment. Our results complement the T cell-centric findings of the NIVO-RELA trial correlative study (25. Andrews *et al.*, 2024; 26. Cillo *et al.*, 2024) and reveal that these two combination ICB regimens also differ in their capacity to induce B cell adaptive immune responses, implicating distinct mechanisms of adaptive immune activation. While our cohort is modest in size, particularly the NIVO-RELA subcohort, it is within the range of comparative neoadjuvant ICB correlative studies (24,27,57); whether the more exuberant TLS and germinal center response translates into more durable outcomes, and whether this differential immune activation affects ICB-induced toxicity, remain important questions for future large-scale studies. Nevertheless, GCSCAN represents a significant advance in computational pathology for TLS quantification, uniquely tailored to the biological complexity of these structures and broadly applicable across imaging and biomarker platforms, fulfilling the recognized need for standardized, molecularly grounded quantification tools to advance TLS-based biomarker development and patient stratification in immunotherapy.

This purpose-built approach reflects a broader principle of our study: each analytical challenge motivated a tailored computational solution when existing general-purpose tools proved insufficient. The same principle guided the development of PathNet-TLS. Despite the availability of HookNet-TLS, benchmarking in our neoadjuvant ICB cohort revealed suboptimal performance for both HookNet-TLS and histopathology foundation models, which incorrectly labeled large regions of immune-enriched tumor bed as confluent TLS and often failed to resolve individual germinal centers. A fundamental limitation is that these models are predominantly trained on treatment-agnostic archival datasets, whose manual annotations are inherently inaccurate in the immune-enriched microenvironments of ICB response.

Uniquely positioned with a paired multimodal dataset of matched H&E, MERFISH, mIF, and same-slide IHC whole-slide images from immunotherapy-treated tumor beds, we generated a molecularly validated ground truth for TLS and germinal center structures. Critically, precise single-cell delineation by GCSCAN replaced manual annotation entirely, yielding training data of unprecedented molecular precision. Trained on 993 mature TLS from co-registered mIF images and 1,170 germinal centers from same-slide Ki67/CD21 IHC, PathNet-TLS replaced subjective visual estimation with objective, molecularly validated boundaries and achieved accurate, scalable TLS and germinal center detection from routine H&E. It reproduced the regimen-specific differences identified by GCSCAN, establishing proof-of-concept for future neoadjuvant trial correlative studies. As the FDA has recognized pathologic response as a surrogate endpoint for accelerated approval across multiple cancer types, the development and validation of computational tools for neoadjuvant response assessment are of direct regulatory relevance.

Beyond the structural quantification of TLS and germinal centers, elucidating the spatial molecular interactions underlying ICB response is critically important to understanding how these organized immune hubs are assembled and sustained through spatially resolved intercellular signaling. In particular, while prior studies identified TCF7+ stem-like T cells as ICB mediators and TLS enrichment in neoadjuvant ICB response, their spatial localization and microenvironmental context in situ remained poorly defined. Our single-cell cellular neighborhood analysis resolved the spatial assembly of distinct T cell, germinal center/B cell, and myeloid zones and localized TCF7+ stem-like T cells within a T cell zone co-enriched with CCL19/CCL21+ fibroblasts and non-germinal center B cells.

Similar to GCSCAN and PathNet-TLS, SCIRA was developed to address the fundamental limitations of existing cell-cell communication algorithms, which either failed at whole-slide scale due to prohibitive computational demands, operated at cell group rather than single-cell level, or required restriction to predefined immune hub regions (22), precluding unbiased pairwise receptor-ligand quantification across millions of cells in whole-slide images. Applying SCIRA across the full MERFISH whole-slide and TMA datasets at single-cell pairwise resolution, we identified zone-specific chemokine receptor-ligand interactions preferentially enriched within the T cell, germinal center/B cell, and myeloid zones of the neoadjuvant ICB response microenvironment, consistent with and extending the cellular neighborhood organization identified by CN analysis.

Most strikingly, SCIRA identified CCL19/CCL21+ fibroblasts as a stromal signal sender to TCF7+ stem-like CD8 and CD4 T cells via the CCR7-CCL19/CCL21 axis, alongside CD40-CD40LG interactions between LAMP3+ dendritic cells and TCF7+ stem-like CD4 T cells, extending prior observations of CCL19+ fibroblast-immune cell interactions to the specific context of stem-like T cell niche organization in the neoadjuvant ICB response microenvironment. SCIRA also recovered the canonical germinal-center circuits (Tfh–GC B via CD40-CD40LG and ICOS-ICOSLG; FDC–GC B via CXCL13-CXCR5), confirming the tool’s fidelity to known biology. Together, these analyses implicate stromal remodeling as a candidate determinant of stem-like T cell niche organization and a potential therapeutic target for strategies aimed at promoting or sustaining stem-like T cell niches in the neoadjuvant ICB setting.

Beyond intercellular signaling, the three-dimensional organization of TLS and germinal center structures represents an entirely unexplored dimension of the neoadjuvant ICB response microenvironment. Leveraging open-top light-sheet imaging and CODA-based 3D reconstruction, we generated the first three-dimensional histologic atlas of the melanoma neoadjuvant ICB tumor ecosystem, revealing striking morphologic heterogeneity across Z-planes and frequently interconnected GC-TLS tunnels that appeared spatially distinct at the 2D level. We further demonstrated that post-OTLS samples can be reverse-cleared for clinical-grade H&E and IHC, establishing a workflow for integrating 3D morphologic imaging with standard immunophenotyping in future studies. These findings raise a fundamental question: is current single-section 2D response scoring sufficient to capture the true extent of the immune response and morphogenic signaling within the treated tumor bed, or does the interconnected three-dimensional architecture of TLS and germinal centers render standard INMC-based assessment fundamentally incomplete? While 3D spatial profiling at clinical scale is not yet feasible, these proof-of-concept observations motivate investigation of 3D imaging and computational pathology protocols as a step toward capturing three-dimensional immune architecture in neoadjuvant response assessment.

In sum, this study represents the most comprehensive single-cell resolution interrogation of the neoadjuvant ICB tumor microenvironment assembled to date, defining the multicellular architecture, intercellular signaling, and immune and stromal networks underlying ICB response and resistance, and extending this into the third dimension through the first three-dimensional histologic atlas of the melanoma neoadjuvant ICB tumor ecosystem. The discovery-to-tool paradigm established here, from MERFISH-informed single-cell discovery through molecularly validated GCSCAN-based quantification to clinically deployable PathNet-TLS H&E-based detection, provides a generalizable blueprint for translating high-dimensional spatial discoveries into universally accessible computational pathology tools, with particular relevance as neoadjuvant immunotherapy expands across cancer types where spatially resolved biomarkers and purpose-built tools remain a critical unmet need. Prospective studies are well-positioned to test whether GCSCAN-based quantification improves pathologic response prediction beyond current INMC visual estimation, and whether PathNet-TLS H&E scoring can serve as a validated surrogate endpoint for response assessment and patient stratification in clinical trials. As mounting evidence establishes TLS as both compelling prognostic biomarkers and promising therapeutic targets in cancer immunotherapy, the single-cell molecular framework, purpose-built computational tools, and neoadjuvant ICB datasets established here provide the foundation for advancing this frontier with spatial precision and translational impact.

## DATA AVAILABILITY

Data generated in this study, including spatial transcriptomics data, raw and processed FFPE single-cell data, and processed mIF data, will be deposited in the Gene Expression Omnibus (GEO) and Zenodo and made publicly available upon publication of the peer-reviewed article. Raw H&E images collected in-house are subject to institutional intellectual property and IRB restrictions.

## CODE AVAILABILITY

All data processing and analysis code will be made available via GitHub and Zenodo upon publication of the peer-reviewed article. The deposited code includes scripts for data processing and for reproducing the main figures and supplementary tables.

## ACKNOWLEDGMENTS

This work was supported in part by the H. Lee Moffitt Cancer Center & Research Institute Total Cancer Care® Research Protocol, Tissue Core, Molecular Genomics Core, and Advanced Analytical and Digital Laboratory, which were supported by the Cancer Center Support Grant (CCSG) P30-CA076292 to H. Lee Moffitt Cancer Center. This research was also supported by generous philanthropic contributions to Moffitt Foundation and Department of Cutaneous Oncology research support fund to PLC. JHC was supported by The Jackson Laboratory Cancer Center Support Grant P30CA034196. This work was also funded, in part, by the Dr. Miriam and Sheldon G. Adelson Medical Research Foundation (to J.J.M). JM has support to the institution from Merck and TuHURA Biosciences. We are grateful to the patients and their families for contributing to this work. We also thank Dr. Jim Mulé and Dr. Patrick Hwu’s lab members for their constructive feedback. Schematic illustrations in this study were created with BioRender.com.

## AUTHOR CONTRIBUTIONS

Conceptualization: ZL, NPR, JHC, PLC; Methodology: ZL, XS, WSC, SD, NPR, JH, LH, CC, JAB, ZS, CMS, NL-B, AA, JVN, JOJ, JHC, PLC; Investigation: ZL, XS, WSC, SD, NPR, CMS, JHC, PLC; Writing – original draft: ZL, NPR, JHC, PLC; Writing – review & editing: ZL, XS, JLM, VKS, JM, PH, JJM, JHC, PLC; Funding acquisition: PLC; Resources: JAB, ZS, CMS, JOJ, DM, SJY, NPR, JLM, VKS, JHC, PLC; Supervision: JHC, PLC

## DECLARATION OF GENERATIVE AI AND AI-ASSISTED TECHNOLOGIES IN THE WRITING PROCESS

During the preparation of this work, the authors used Claude to improve the readability and language of the work. After using this tool, the authors reviewed and edited the content as needed and take full responsibility for the content of the publication.

## SUPPLEMENTAL INFORMATION

Document S1. Supplemental Figures S1–S10

Table S1. Cohort and tissue amples

Table S2. Clinical demographics

Table S3. MERFISH gene panel

Table S4. MERFISH cell-typing markers

Table S5. RL pairs in MERFISH gene panel

Table S6. MERFISH data, tissue composition by cell types (related to Figures 2E and S5)

Table S7. 9-plex mIF data, tissue composition by cell types (related to Figures 2I and 2K)

Table S8. SCIRA results, aggregated receiver-sender R-L interaction strength, MERFISH data (related to Figure 4C)

Table S9. GCSCAN results, aggregated at patient level for all MERFISH and 4-plex mIF data (related to Figure 5)

Table S10. PathNet-TLS mTLS and GC quantification (related to Figure 6)

Video S1-4. 3D imaging and rendering of responder tissue architecture, interconnected TLSs, and high endothelial venules inside an individual germinal center

## METHODS

### Experimental Model And Study Participant Details

We retrospectively assembled a cohort of a total of 87 patients with stage III metastatic melanoma, including 60 neoadjuvant ICB treated (25 IPI NIVO, 21 NIVO RELA, and 14 NIVO/PEMBRO monotherapy) and 27 treatment naïve (TxN) (Figure 1A; Table S1). All biospecimens in this study were performed on retrospective FFPE tissues collected as part of routine patient care. All tumor bed surgical specimens were collected at time of curative intent surgery and included digitally scanned H&E images. A total of 31 complete response (CR), 7 partial response (PR), 22 non-response (NR), 4 uninvolved LN and 27 TxN tumor bed FFPE blocks were collected for single-cell/spatial –omics, 3D open-top light-sheet (OTLS) imaging and 3D H&E image reconstruction by CODA.

### Digital Pathology

All H&E and immunohistochemistry slides were digitally scanned on a Leica Aperio GT450 scanner at 40× magnification. To assess pathologic response in the neoadjuvant ICB samples, resected surgical specimens were scored according to the International Neoadjuvant Melanoma Consortium guidelines (14), using the percentage of residual viable tumor (RVT) together with fibroinflammatory stroma, necrosis, and pigmented macrophages. Response categories were defined as complete response (CR; 0% RVT), pathologic major response (pMR; ≤10% RVT), partial response (PR; >10% and ≤50% RVT), and non-response (NR; >50% RVT). For this study, pMR patients were considered as CR, and both CR and PR were considered as responders. Tumor bed areas were annotated on H&E based on evidence of previous tumor involvement.

### FFPE scRNA seq (Flex Fixed RNA Profiling, four plex)

#### FFPE tissue processing and data acquisition

Single-cell RNA-sequencing from fixed tissue was performed by the Moffitt Cancer Center Molecular Genomics Core using the 10X Genomics Chromium Single Cell Gene Expression Flex Kit (10X Genomics, Inc., Pleasanton, CA). Formalin-fixed paraffin embedded tissues were dissociated by the Moffitt Cancer Center Tissue Core according to the 10X Genomics protocol. Two 50 μm FFPE tissue sections were deparaffinized through three sequential 10-minute xylene incubations, followed by a graded ethanol rehydration series (100%, 100%, 70%, and 50% ethanol), a water wash, and PBS equilibration on ice. After deparaffinization, the tissue is incubated in a Liberase TH-based Dissociation Enzyme Mix and processed on a gentleMACS Octo Dissociator using the manufacturer’s FFPE program for 48 minutes to enzymatically and mechanically dissociate the tissue into single cells. The dissociated material was filtered through a 30μm strainer to remove debris and the cells pelleted by centrifugation. Cells were resuspended in Quenching Buffer and counted using the Nexcelom Cellometer K2 (Nexcelom Bioscience, Lawrence, MA). Up to 2 million cells of each sample were hybridized overnight with a separately barcoded pool of Human WTA mRNA detection probe pairs, and after preparing fourplex sample pools and washing, the pool was encapsulated using the 10X Genomics Chromium X Single Cell Controller at a concentration of four thousand cells per microliter to target 10,000 cells per sample. Briefly, the single cells, reagents, and 10x Genomics gel beads were encapsulated into individual nanoliter-sized Gelbeads in Emulsion (GEMs) and then following ligation of mRNA-bound probes, pre-amplification was performed inside each droplet. The indexed DNA libraries were then completed in a single bulk reaction and approximately 20,000 sequencing reads per cell were generated on the Illumina NovaSeq 6000 instrument (Illumina, Inc., San Diego, CA). Demultiplexing, barcode processing, alignment, and gene counting was performed using the 10X Genomics CellRanger software.

#### Preprocessing

Raw sequencing reads were demultiplexed and converted to FASTQ format using Cell Ranger mkfastq (v7.1, 10X Genomics). The resulting FASTQ files were then processed with cellranger multi. Specifically, mRNA sequencing reads were aligned against GRCh38 human transcriptome, cellular barcodes were assigned, and unique molecular identifiers (UMIs) were counted to generate a gene-by-cell count matrix. Downstream analysis was performed in Seurat (v4.2.2) (58).

#### Feature and cell filtering

Cells were removed if they met any of the following criteria: i) ≤ 500 genes detected, ii) ≤ 500 transcripts, iii) ≥ 10% mitochondrial RNA content, or iv) 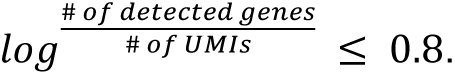 To identify and remove doublets, cells were further excluded if they were predicted as a doublet by Scrublet or contained an abnormally high number of transcripts. The upper threshold for transcript count was defined as the median plus four times the median absolute deviation (MAD). In addition, genes detected in less than ten cells were excluded.

#### Melanoma cell identification

InferCNV (v1.10.1) (59) was used to identify melanoma cells from CD45⁻ cells (defined as cells with at least one sequencing read mapped to PTPRC). CD45⁻ cells from lymph node (LN) control samples were used as a reference, against which chromosomal CNV patterns were compared in CD45⁻ cells from each melanoma sample. A cutoff of 0.1 was applied, and denoising was enabled. The cluster_by_groups parameter was turned off to allow unsupervised hierarchical clustering based on CNV patterns, under the hypothesis that benign and malignant cells would segregate into distinct histogram clusters. To better highlight benign cells, half of the LN CD45⁻ cells were randomly selected and placed in the “observation” group as controls. Cells that clustered separately from LN spike-in cells and exhibited CNV patterns underwent additional selection based on melanoma markers. Final melanoma cells were identified as SOX10⁺/MLANA⁺ cells with malignant CNV patterns.

#### Non-melanoma cell analysis

For each sample, SCTransform (v2) was applied while regressing out potential biases introduced by mitochondrial RNA content. To correct for batch effects across samples, the Seurat integration workflow was performed using the top 5,000 highly variable genes. Principal component analysis (PCA) was performed on the integrated dataset to reduce dimensionality. The first 30 principal components (PCs) were used to construct a shared nearest neighbor (SNN) graph, which served as the basis for clustering cells using the Louvain algorithm at multiple resolutions. To visualize the data in two dimensions, Uniform Manifold Approximation and Projection (UMAP) was generated using the top 40 PCs. To characterize and annotate the identified clusters, orthogonal approaches were applied: i) Assessment of established cell type-defining markers by examining their gene expression patterns across cell clusters; ii) Differential expression analysis was performed using FindAllMarkers function, comparing each cluster to all others; iii) Cross-referencing with bioinformatically predicted cell types based on both public datasets (SingleR, Monaco immune data) and an in-house single-cell dataset (Azimuth, melanoma samples). Cell type for each cell cluster was manually reviewed and assigned. Benign non-immune cells were selected from those CD45-cells lacking malignant CNV patterns (see Melanoma Cell Identification) by further excluding cells that expressed SOX10 or MLANA. Following data processing, various non-immune cell types were identified, and T and B cells affected by drop-out issues were recovered. Raw count data of these cells were then jointly processed with the immune compartment of TME, defined as CD45⁺ SOX10⁻ MLANA⁻ cells, using the same approach. Cell types for each cluster were manually assigned and visualized using UMAP.

### MERFISH dataset generation and processing

#### Panel design and sample preparation

To profile immune cell types, cell states, and key interactions in situ, we curated a panel of 305 genes based on prior literature and submitted it to Vizgen for MERFISH 2.0 chemistry gene panel design (34) (https://portal.vizgen.com/). Human samples were prepared as FFPE blocks and sectioned at 5 µm onto MERFISH Standard Slide V2.0 (Vizgen, 20400117). Sample preparation followed Vizgen’s MERFISH 2.0 Sample Preparation User Guide for Sectioned Tissue Samples (Vizgen, 91600132) and instrumentation followed the MERFISH Instrument Guide (Vizgen, 9160001); study-specific reagents and deviations are summarized below.

The tissue sections were deparaffinized by Deparaffinization Buffer (Vizgen 20300112) at 55°C for 5 minutes twice, and washed with 100% ethanol three times, each 2 minutes, followed by a 2 minute 90% ethanol and a 2 minute 70% ethanol rehydration step. The rehydrated tissue sections were incubated with Decrosslinking Buffer (Vizgen 20300115) at 90°C for 15 min and cooled on bench for 5 minutes. The decrosslinked tissue sections were then stained for cell boundary using Vizgen’s Cell Boundary Kit (10400118) following Vizgen’s MERFISH 2.0 Sample Preparation User Guide for Sectioned Tissue Samples (Vizgen, 91600132). After post fixing the sample with 4% PFA for 15minutes, the tissue sections were incubated with Conditioning Buffer (Vizgen 20300116) supplemented with RNase inhibitor at room temperature for 15 min, and then Pre-Anchoring Reaction Buffer (Vizgen 20300113) overnight at 37°C in a humidified chamber. Following anchoring pretreatment, tissue slices were then briefly washed with Sample Prep Wash Buffer (Vizgen, 20300001), and Formamide Wash Buffer for 15 min at 37°C (Vizgen, 20300002), and then Anchoring Buffer (PN 20300117) at 37°C for 2 hours. Afterwards, the sample was washed with Sample Prep Wash Buffer briefly and then gel embedded and cleared using the MERFISH Tissue Sample Prep Kit (Vizgen, 10400194) overnight at 47°C. After tissue clearing, the sample was treated with MERFISH Photobleacher (Vizgen 10100003) for 3 h, washed with Formamide Wash Buffer at 37°C for 30 min, and then incubated with MERFISH Gene Panel Mix V2.0 at 47°C overnight. The sample was then incubated with Enhancer (Vizgen, 30300491) at 37°C overnight, and washed with Enhancer Wash Buffer (Vizgen, 20300192) at 37°C for 20 min, twice. The sample was stained with DAPI and Poly T Reagent V2.0 for 15 min at room temperature, washed for 10 min with 5 mL of Formamide Wash Buffer, and then imaged on the MERFISH M1 system (Vizgen 10000001) using the MERFISH Standard 500 Gene Imaging Kit V 2.0 (Vizgen, 10400169). A fully detailed, step-by-step instructions on the MERFISH sample prep can be found in Vizgen’s MERFISH 2.0 Sample Preparation User Guide for Sectioned Tissue Samples (Vizgen, 91600132). The full instrumentation protocol can be found in MERFISH Instrument Guide (Vizgen, 9160001).

#### Cell segmentation

To recover single-cell boundaries with high fidelity, we used a two-step Cellpose (60) and Baysor (61) approach. After image acquisition, the MERFISH data was processed by Vizgen, and the Cellpose algorithm was used to perform cell segmentation. Cellpose, as implemented in the Vizgen Post-processing Tool (VPT), was applied to DAPI and cell-boundary staining (Cell Bound 3 from Vizgen’s Cell Boundary Kit), with tissues tiled and stitched within VPT. Using the Cellpose result as a prior, Baysor (v0.6.2) segmentation was performed with default parameters and a prior confidence of 0.8. For Baysor, tissues were tiled into 2800 × 2800 tiles with 300-pixel overlap per side and stitched back together, retaining only cells in the center 2500 × 2500 area. Single-cell expression matrices were generated by aggregating detected transcripts by Baysor cell ID, weighted by assignment confidence.

#### Identification of individual cores in TMA arrays

To analyze TMA cores independently, cores in the three TMAs were detected and labeled using a foreground-detection approach (see code for implementation details). Segmented cell polygons were rasterized into a binary cell mask and dilated by 50 pixels (binary_dilation(), scikit-image v0.22.0) to close inter-cell gaps. Cores were detected as foreground connected components (label(), scikit-image), and a minimum-area threshold computed by Otsu’s method was used to filter out small false positives. To prevent adjacent cores from merging during dilation, we compared the maximum and median bounding-box areas and reduced the dilation radius whenever the maximum-to-median ratio exceeded 1.5, which indicated that neighboring cores had been joined into one region. Detected cores were then aligned onto a grid using an iterative breadth-first search: starting from a random core, the algorithm moved along the four cardinal directions using 1.25 times the median core height or width as the estimated center-to-center distance, snapping to a neighboring core center when one was found and marking the position as a skipped core otherwise. The labeled grid was matched to the H&E core-labeling system or serially numbered.

#### Quality control and normalization

To avoid cross-batch effects, each MERFISH whole-slide image (WSI) and TMA slide was processed separately. Cells with fewer than 10 transcripts, or fewer than 5 unique genes with at least 1 transcript each, were removed as low-quality. Three of the 30 treatment-naïve TMA cores had more than 60% of cells removed and were excluded. After quality control, 4,837,032 cells were retained across all samples. Read counts were normalized by the median of non-zero read counts for each gene across all cells in each batch and log-transformed (log(1 + *x*)). For samples with tumor-bed annotation, a tumor-bed mask was generated by overlaying the annotated H&E on the MERFISH tissue image and tracing the annotation.

#### Clustering and cell type annotation

To account for patient-specific and technical batch effects, each imaging batch was clustered and annotated separately. The gene-wise normalized cell-by-gene matrix was reduced to 50 components by non-negative matrix factorization (NMF, scikit-learn v1.4.1, Multiplicative Update solver). A nearest-neighbor graph (15 neighbors, cosine distance) was built with NearestNeighbors() (scikit-learn) and igraph v0.11.5, and clustered by Leiden clustering (46) (resolution 2, find_partition(), leidenalg v0.10.2). First-round clusters were annotated into major cell types using known markers (Table S4). For each major type, a second round of graph construction and Leiden clustering refined the cell typing, annotated using known markers and top-ranked genes (ScanPy v1.9.8). Clusters lacking expression of all known markers were labeled “undefined.” For clusters with a multi-lineage expression pattern, a non-negative Lasso regression (scikit-learn v1.4.1, alpha = 0.1) decomposed the expression into a linear combination of the average expression of other single-cell-type clusters within the same sample. Components with cell-type-specific weights > 0.5 were reconstructed into their single-cell-type identities, remainders were excluded, and clusters that could not be explained as such a combination were labeled “undefined.”

#### Cellular Neighborhood (CN) identification and analysis

To characterize each cell’s local tissue environment, we computed a CN expression profile for every cell as the mean expression across its 10 nearest neighbors, equivalent to a graph convolution with uniform weights. CN profiles were reduced to 50 NMF components and clustered by Leiden clustering (resolution 2), using the same NMF and Leiden settings described for single-cell clustering. The number of CN Leiden clusters was reduced to 9 by agglomerative hierarchical clustering (HC, hierarchy module, SciPy v1.12.0) on the mean features of each CN Leiden cluster (Euclidean distance, Ward’s method). HC clusters were manually annotated by their spatial distribution and dominant cell types. The HC clusters and hierarchical trees were inspected for potential necessary merging or splitting to yield meaningful CN groups.

To characterize the cell-cell colocalization behaviors in relation to the local spatial niche, we calculated neighbor enrichment for each cell type occurring within each CN group (here we use “CN” to refer to the overall CN group from clustering, not individual spatial instances within the cluster). For each cell, the neighboring cell-type frequency was computed as the frequency of each cell type within a 50-pixel radius, using a denominator of five for cells with fewer than five neighbors. For a given center cell type, these frequencies were averaged across instances within the CN to give an average neighboring cell-type frequency vector. To assess significance and reveal CN-specific patterns, a permutation test randomly sampled the same number of center-type cells within the current CN from all CNs, and z-scores were computed against the distribution from 1,000 random samplings.

#### Spatial Cellular Interaction and Receptor Activation (SCIRA)

The purpose of SCIRA (62) is to quantify spatial receptor-ligand interactions for each single cell. For a given receptor-ligand (R-L) pair, the signal received by a cell from its neighbors was quantified with a simplified mass-action kinetics model:

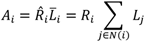

where *A_i_*, is the strength of receptor activation by the corresponding ligand in cell *i*, R̂i is the receptor expression level in that cell, and *L_i_* is the total ligand expressed by its neighboring cells *N*(*i*). The neighborhood *N*(*i*) comprises all cells whose centers lie within a distance threshold of the center of cell *i*, excluding cell *i* itself. We used a radius of 15 µm (15 pixels), selected to account for surface-bound and short-range diffusible ligands; this radius was empirically verified to capture diffusible ligand–receptor interactions in our data. The total neighbor ligand level *L̄_i_*, representing access to the ligand in the local environment, was computed as the sum of ligand expression across neighboring cells, optionally distance-weighted for longer-range interactions.

To aggregate these single-cell measurements to the cell-type level, the following quantities were computed for each R–L pair.

First, the receiver-centric receptor activation score Ā*_r_* is the mean single-cell receptor activation across cells of receiver type *r*:

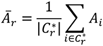

where *C_r_*^∗^ is the set of cells of type *r* having at least one neighbor.

Second, *Î*_s→r,_ the share of contribution of sender cell type *s* to the signal received by receiver cell type *r* is obtained by normalizing each sender type’s contribution across all contributing sender types for the receiver type:

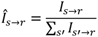

where *I_s_*_→*r*_ is the unnormalized contribution of sender cell type *s* to the total signal received by receiver cell type *r*. It is calculated as the ligand expression of each signal-sending cell multiplied by the number of receiver-type cells in that cell’s neighborhood, averaged over signal-sending cells of type *s*:

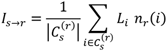

where *L_i_* is the ligand expression in signal-sending cell *i*, *n_r_*(*i*) is the number of receiver-type cells in the neighborhood of cell *i*, and *C_s_*^(*r*)^ is the set of cells of sender type *s* having at least one neighbor of receiver type *r*. Sender cells with no receiver-type neighbor are excluded from the average calculation, so that an abundant sender type is not penalized by a low proportion of engaged cells.

Finally, we define *S_s_*_→*r*_, the interaction strength between a sender-receiver cell-type pair for a given R-L pair, as the product of the mean receiver activation score and the share of signal contribution of the sender cell type:

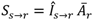

These aggregated quantities are defined for a single R-L pair. To obtain values that are comparable across R–L pairs and across cell-type pairs when assembling the results across all R-L pairs for visualization purposes, each R–L pair was rescaled by its maximum over all sender-receiver cell-type pairs:

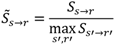

SCIRA was run for MERFISH WSI and TMA data using the 45 R-L pairs in the gene panel (Table S5). To avoid numerical instability, TCF7− CD4 T cells, cDC, and pDC cells were excluded from the results due to rarity and uneven presence across the samples.

For benchmarking, we ran COMMOT (43) (v0.0.3) on individual MERFISH TMA cores with the same radius and the same curated receptor-ligand pair list as SCIRA. COMMOT could not be run on some whole-tissue sections because the initial pairwise distance matrix required more than 700 GiB of memory, exceeding the maximum RAM available on our high-performance computing clusters. TMA core results were used to calculate the concordance between the two methods.

To assess agreement between SCIRA and COMMOT, per-cell-type receptor activation strength values for each receiver cell type for TMA core were matched for each R-L pair. Spearman’s rank correlation coefficient was computed for each pair for each core across cell types. To summarize concordance across cores for each L–R pair, we calculated the median Spearman ρ and interquartile range (IQR; 25th–75th percentile) across cores. Results are presented as a forest plot, with L–R pairs ordered by median correlation and error bars denoting the IQR.

### Vectra Polaris Multiplexed Immunofluorescence (mIF) data generation and analysis

#### Data generation

Formalin-fixed and paraffin-embedded (FFPE) tissue samples were immunostained using the AKOYA Biosciences OPAL™ 7-Color Automation Immunohistochemistry (IHC) kit (Waltham, MA) on the BOND RX autostainer (Leica Biosystems, Vista, CA). The OPAL 7-color kit uses tyramide signal amplification (TSA)-conjugated to individual fluorophores to detect various targets within the multiplex assay.

Sections were baked at 65 °C for one hour then transferred to the BOND RX (Leica Biosystems). All subsequent steps (e.g., deparaffinization, antigen retrieval) were performed using an automated OPAL IHC procedure (AKOYA). OPAL staining of each antigen occurred as follows: heat-induced epitope retrieval (HIER) was achieved with HIER-EDTA pH 9.0 buffer for 20 min at 95 °C before the slides were blocked with AKOYA blocking buffer for 10 min. Slides were then incubated with the primary antibody, Ki67 (Abcam, polyclonal, 1:100), at RT for 60 min, followed by OPAL HRP polymer and one of the OPAL fluorophores during the final TSA step. Individual antibody complexes were stripped after each round of antigen detection. This was repeated using the following antibodies: BCL6 (Abcam, polyclonal, HIER-EDTA pH 9.0, 1:100), PAX5 (Abcam, EPR3730(2), HIER-EDTA pH 9.0, 1:100), CD21 (Abcam, EP3093, HIER-EDTA pH 9.0, 1:400), and CD3 (Dako, polyclonal, HIER-EDTA pH 9.0, 1:300). The following antibodies were used on separate mIF panels: CD45 (Abcam, rabbit polyclonal, HIER-EDTA pH 9.0, 1:[CD45 dilution: 1:100]), LAMP3 (Invitrogen, rabbit polyclonal, HIER-EDTA pH 9.0, 1:100), TCF1/7 (CST, C63D9, HIER-EDTA pH 9.0, 1:100), CD38 (CST, E7Z8C, HIER-EDTA pH 9.0, 1:300), and CD8 (Dako, C8/144B, HIER-EDTA pH 9.0, 1:50). After the final stripping step, DAPI counterstain was applied to the multiplexed slide, which was then removed from the BOND RX for coverslipping with ProLong Diamond Antifade Mountant (ThermoFisher Scientific). All slides were imaged with the AKOYA PhenoImager HT Imaging System.

A total of 54 neoadjuvant-treated patients (28 CR, 6 PR, 20 NR) were profiled by Vectra multiplexed immunofluorescence (mIF). First, a 9-plex panel (DAPI, CD3, CD8, TCF1/7, CD38, CD45, LAMP3, PAX5, CD21) was applied to 29 patients (14 CR, 4 PR, 11 NR) for tissue composition analysis. A 4-plex panel (DAPI, CD21, Ki67, BCL6) was then applied across the entire 54-patient cohort for TLS and germinal center (GC) analysis, and quantitative TLS/GC measurements were derived from these 4-plex images. Additionally, to resolve B-cell and T-cell distributions within the 4-plex panel, the 25 patients profiled with only the 4-plex panel had their sections stained for PAX5 and CD3 and re-imaged.

#### Image processing

To suppress noise and autofluorescence prior to segmentation, each 9-plex channel was denoised with a 2-component Gaussian Mixture Model (GMM) trained on a subsample of 1 × 10⁶ pixels, weighting each pixel by its probability of belonging to the higher-mean component. The autofluorescence channel was subtracted from CD38, and CD8 pixel intensity was capped by CD3 intensity. Each channel was rescaled to 0–1, with 1 corresponding to the 90th percentile (CD45) or 99th percentile (other markers) of non-zero pixels in the raw image. The first round 4-plex images were denoised using the same GMM-based method, and rescaled to 0–1 with the 99^th^ percentiles. The second round 4-plex images were normalized per tile during the cell segmentation step because of the presence of uneven staining across the whole section.

#### Cell segmentation

Cell segmentation was performed using DeepCell (v0.12.9) Mesmer whole-cell segmentation algorithm (63) with the pretrained Multiplex Segmentation-9 weights at image mpp = 0.5. For the 9-plex images, the maximum projection of DAPI and Ki67 channels was used as the nuclear channel, and the maximum projection of CD3, CD45, CD8 and CD21 channels was used as the membrane channel. For the 4-plex images, the DAPI channel was used as the nuclear channel and the CD21 channel was used as the membrane channel. The whole section images were first tiled into 4000 x 4000 tiles with 1000 pixel overlapping on each side before segmentation and stitched back together keeping only the cells in the center 3000 x 3000 area after segmentation. Whole-cell average intensity of each marker inside each segmented cell was calculated as the single-cell protein expression matrices. An above-upper-quantile average was also calculated, which was the mean intensity of the brightest 25% of pixels inside each cell.

#### Clustering and cell type annotation

For 9-plex images, the single-cell expression matrix was clustered by agglomerative hierarchical clustering (SciPy v1.12.0) into 15 clusters, annotated by average expression. For 4-plex images, a threshold of 0.1 defined positivity for CD21 and Ki67, and cells were annotated as CD21⁺Ki67⁺, CD21⁺Ki67⁻, or CD21⁻. CD21 uses whole-cell mean, and Ki67 used above-upper-quantile mean. The thresholds and features were chosen by inspection of per-marker-intensity histograms. Images with abnormal staining were corrected by visual inspection.

#### Image registration and tumor-bed annotation transfer

To transfer tumor-bed (TB) and outside-tumor-bed (OTB) annotations from H&E in QuPath to the Vectra mIF images, we registered the H&E hematoxylin channel to the mIF DAPI channel using the top 25% of 15,000 Oriented FAST and Rotated BRIEF (ORB) features matched by the Brute-Force matcher (OpenCV-Python v4.9.0). The resulting affine transformation was applied to the TB annotation polygon (Polygon instances, shapely v2.0.3) in the GeoJSON exported from QuPath (v0.4.4). Registration was visually inspected per sample, and annotations were transferred manually for poorly registered samples.

### Graph-based Cellular Spatial Clustering Against Noise (GCSCAN) Algorithm

The purpose of GCSCAN was to detect spatially clustered cellular structures such as TLS and follicles. We provide a high-level description here and refer to the GitHub repository for implementation details (64).

To represent spatial relationships, we first constructed a weighted undirected graph (a Graph instance in NetworkX v3.2) in which vertices are cells and edges reflect spatial adjacency. For MERFISH samples, adjacency was computed from the minimum distance between dots on the boundaries of two neighboring cells, connecting only cell pairs with a boundary distance below 5 µm (5 pixels). For Vectra mIF samples, Delaunay triangulation (SciPy v1.12.0) was used with a maximum center-to-center distance of 25 pixels.

To separate follicular from non-follicular cells, a node-level “involvement score” was computed for each cell as the fraction of TLS/follicle-related cell types among its neighbors, so that cells inside structures scored high and those outside scored low. This step is critical for distinguishing B cells within and outside follicles, which is the main source of false positives (noises). TLS/follicle-related cells were B-GC, B-non-GC, Tfh, and FDC for MERFISH samples and all CD21⁺ cells for mIF samples. Each edge was weighted by the product of the involvement scores of its two vertices.

To delineate structures, a modified graph-based watershed algorithm was used. This started from high-involvement nodes and expanded along the graph until reaching low-weight edges. Adjacent structures loosely connected by few cells were then separated by Spectral Graph Partitioning (SpectralClustering(), scikit-learn v1.4.1), dividing subgraphs whose normalized cut cost fell below the predetermined threshold of 0.005. Each subgraph underwent graph-based dilation and erosion to refine its shape, and structures with more than 200 cells were retained for counting and statistical comparison. Follicle and GC contours were approximated by the convex hull of all cells on the alpha-shape boundary, with the alpha value estimated from the mean nearest-neighbor distance. Each sample was visually inspected to remove false positives arising from staining or tissue artifacts.

### TLS and GC comparisons

For mIF comparisons, CD21⁺ follicle cells were CD21⁺ cells inside any detected structure, and a follicle was scored as GC-positive when at least 15% of its cells were CD21⁺Ki67⁺ and located inside the approximated GC area. Sample-level averages were grouped by response and treatment or by response alone. P-values for comparisons were assessed using the Mann-Whitney U test. For tumor-bed-annotated samples, only cells and structures inside the annotated tumor bed were included.

For benchmarking, we ran HDBSCAN on all B-GC, B-non-GC, Tfh, and FDC cells in the same MERFISH samples (min_cluster_size = 20, min_samples = 30, cluster_selection_epsilon = 1), with contours identified by the same alpha-shape convex-hull approach. TLS cell percentages were calculated for each detected mTLS as the number of cells of each TLS-specific cell type (GC B cells, non-GC B cells, FDC, and Tfh) divided by the total number of cells inside the mTLS.

### Survival analysis with GCSCAN

Overall survival (OS) was defined as time from neoadjuvant immune checkpoint blockade (NICB) treatment start date to death from any cause or last documented follow-up for surviving patients. Disease-free survival (DFS) was defined as time from NICB treatment start date to pathologic progression of disease (POD) or death, whichever occurred first. Patients without documented progression were censored at last follow-up or death. To avoid instability in Kaplan-Meier estimates and log-rank statistics arising from very small risk sets at late timepoints, follow-up time was censored at a data-driven cutoff, defined per analysis as the earliest timepoint at which any compared group’s number-at-risk fell to ≤3 patients, subject to a hard ceiling of 3 years. This truncation was specified and applied prior to computing both the Kaplan-Meier curves and all associated log-rank statistics. Number-at-risk tables, including cumulative censoring counts, are displayed below each Kaplan-Meier plot at fixed time intervals. Kaplan-Meier estimates were computed in Python using the lifelines package (v 0.27.8). Continuous variables were dichotomized at the cohort median to define comparison groups for survival stratification. Differences in OS/DFS across groups were assessed using the log-rank test.

### TLS detection on H&E images with UNI features

#### Image preprocessing and feature extraction

To generate tile-level representations for downstream TLS detection, WSIs were processed with the STQ pipeline (https://github.com/TheJacksonLaboratory/STQ/, v0.3.0) (65). All images were processed at 40× resolution (0.25 µm/pixel) to preserve fine texture. Stain normalization was performed used the Macenko method with a fixed reference and a 512-pixel patch, tissue regions were identified by intensity masking (background cutoff 210.0), and focus quality was assessed with a pretrained DeepFocus model, all implemented in STQ. Non-overlapping 56 µm tiles (224 × 224 pixels at 0.25 µm/pixel, matching the UNI input size) were generated on a fixed grid, and 1,024-length feature embeddings were extracted per tile with the pretrained UNI model (38).

#### Ground-truth label generation

Ground-truth TLS labels were generated by registering matched H&E and Vectra mIF images and transferring polygons of GCSCAN-detected TLS regions (as described in the PathNet-TLS section, which uses the same 13 representative slides). TLS regions with more than 200 and less than 4,000 cells were retained for training. Each tile was labeled by whether its center fell inside or outside a transferred TLS region. All TLS-positive tiles were kept and twice as many TLS-negative tiles were randomly sampled, yielding 47,469 labeled tiles.

#### Multiscale spatial context embedding

Although the tiles at high resolution preserve fine-scale cellular morphologies, they may lack broader spatial context that are key to distinguishing organized TLS structures from unorganized lymphoid aggregates. To incorporate the spatial context, a radial multiscale feature aggregation strategy was applied to the UNI embeddings. Starting with each tile as the center tile, the feature vector of the center tile was concatenated with the mean feature vectors of neighboring tiles at the expanding frontier at increasing spatial radii. Specifically, first-order neighbors (8 tiles), second-order neighbors (16 tiles), and subsequent neighborhood rings were aggregated up to a maximum of five hops. At each hop, the mean feature vector of all tiles at that hop distance was computed and concatenated to the center tile representation, generating 6144-feature vectors. This resulted in a multiscale representation that captures both local and broader, distance-aware tissue context while maintaining a fixed and computationally efficient embedding length.

#### Downstream classifier

Classification was performed using logistic regression (LR, scikit-learn v1.4.1) and a multilayer perceptron (MLP, TensorFlow v2.16.1). The MLP had two fully connected ReLU hidden layers followed by a sigmoid output, with hidden dimensions (256, 64) for single-tile features and (512, 128) for contextual concatenation. Models were trained with the Adam optimizer and binary cross-entropy loss (learning rate 1 × 10⁻³, weight decay 1 × 10⁻⁵, batch size 256, dropout 0.3, up to 60 epochs), with data split 0.7:0.15:0.15 into training, validation, and test sets. Performance was evaluated by ROC-AUC and PR-AUC during training, and spatial prediction maps and predicted boundaries were visually inspected on both training and held-out samples.

### Immunohistochemistry (IHC)

Immunohistochemistry was performed on automated stainers, and the primary antibodies employed were CD21 (Leica Biosystems, NCL-L-CD21-2G9, 1:100) and Ki67 (Ventana Medical Systems, 790-4286, ready to use). All slides were stained using previously optimized conditions with appropriate positive and negative controls. CD21 was stained on the Leica Bond platform with UltraView Universal Alkaline Phosphatase Red Detection Kit (Ventana Medical Systems). Ki67 was stained on the Ventana BenchMark ULTRA platform with UltraView DAB detection (Ventana Medical Systems). All chromogenic IHC slides were digitally scanned as described above (Digital Pathology).

### PathNet-TLS

The purpose of PathNet-TLS (66) was to identify TLS and germinal centers (GCs) from H&E images. We trained a transformer-based model with a TLS module and a GC module, based on the UNet-like, dual-transformer UDTransNet architecture (53), with the training code modified to incorporate Hugging Face Accelerate (67) for multi-GPU compatibility.

### TLS module

The TLS module was trained using GCSCAN-detected TLS and follicles in Vectra mIF data as ground truth. To transfer labels to H&E, the H&E hematoxylin channel and the mIF DAPI channel were registered to the GCSCAN-detected structures using the top 50% of 15,000 ORB features matched by the Brute-Force matcher (OpenCV-Python v4.9.0), and the TLS/follicle boundary coordinates were transferred onto the H&E images. Registration quality was visually assessed to retain only well-registered samples. A total of 13 representative samples with good registration quality were used to generate TLS ground truth.

Whole-slide H&E images at 10× resolution were divided into 50%-overlapping patches at two tile sizes (448 and 896 pixels) to enable multi-scale learning, with tissue foreground detected from the (red + blue) minus green channel difference. The TLS/follicle annotation was converted to a binary mask and patched identically. Any patch with more than 15% TLS area was treated as positive, with an equal number of negative patches randomly selected. Patches from well-registered samples were split 4:1 into training and validation, training patches were augmented by rotation and flipping, and images were normalized using ImageNet mean and standard deviation, as the convolutional layers were initialized with ImageNet-based ResNet34 weights.

A 5-fold cross-validation ensemble was trained for each tile size (adaptive cosine learning rate starting at 0.0003, batch size 8, up to 500 epochs, early-stopping patience 40). The loss combined a hard DICE term, a BCE term, and a negative DICE term applied only to negative patches to penalize false positives, weighted 1.5:1:0.1. The final prediction was the average predicted probability of the best model per fold, averaged over overlapping patches.

To segment individual structures from the probability mask, the mask was Gaussian-smoothed and binarized at 0.5, processed with binary opening and closing to remove small artifacts, and smoothed again. Regions with intensity > 0.98 were taken as sure foreground and those < 0.5 as sure background. The sure foreground was eroded and distance-transformed for Watershed segmentation, with the distance map smoothed to remove false local maxima. Watershed segmentation then combined the smoothed distance map, local-maxima seeds, and sure-background mask. Morphological operations used OpenCV except the Gaussian filter (scikit-image, v0.22.0), and per-slide counts and areas of detected structures were returned.

### GC module

The GC module was trained using IHC staining on selected slides as ground truth. Matched same-section IHC and H&E were registered using the red channel and the method above. To convert IHC to a bitmap mask, IHC images were Gaussian-smoothed and binarized, then processed with remove_small_holes, MORPH_CLOSE, and remove_small_objects (scikit-image and OpenCV). The IHC and GC bitmaps were tiled into 448 × 448 patches. Training, inference, and segmentation followed the TLS module, except that GC prediction was restricted to TLS areas predicted by the TLS module.

The counts of GCs per slide are irrelevant to the training, but calculated for reporting purposes. It is calculated by segmenting the foreground mask following the remove_small_objects operation using the same watershed-based segmentation method for predicted object segmentation, and filtered by at least 75% overlap with any predicted TLS objects to remove false-positives caused by other Ki67+ proliferative structures.

PathNet-TLS requires no further training for future use. The software takes a histopathology image and returns a GeoJSON file that can be imported into viewers such as QuPath.

### 3D imaging with OTLS microscopy

#### Staining and clearing

FFPE tissues were deparaffinized then washed in 100% ethanol twice for 1 h each to remove any excess xylene. The tissues were then gradually transferred to 100% methanol, followed by delipidation in 70% dichloromethane and 30% Methanol for 8 hours. Following this, the tissues were put through 5 rounds of depigmentation of 2 hours each in a solution of 5% H2O2 in Methanol at 4°C. The tissues were gradually transferred to 100% ethanol then treated in 70% ethanol for 1 h to partially re-hydrate them. Each tissue sample was then placed in an individual 5ml Eppendorf tube (Cat: 14-282-300, Fisher Scientific), stained for 48 hours in 70% ethanol at pH 4 with a 1:200 dilution of Eosin-Y (Cat: 3801615, Leica Biosystems) and a 1:500 dilution of To-PRO™-3 Iodide (Cat: T3605, Thermo-Fisher) at room temperature with gentle agitation. The tissues were then dehydrated twice in 100% ethanol for 2 hours. Finally, the tissues were optically cleared (n = 1.56) by placing them in ethyl cinnamate (Cat: 112372, Sigma-Aldrich) for 8 hours before imaging them with hybrid open-top light-sheet (OTLS) microscopy (35). A custom-machined HIVEX plate (n=1.55) was used as a multi-sample holder (4 tissue samples per holder).

#### Hybrid open-top light-sheet (OTLS) microscopy and data processing

Multi-channel illumination was provided by a four-channel digitally controlled laser package (Skyra, Cobolt Lasers). Entire samples were imaged at 2 µm/pixel resolution with a hybrid open-top light-sheet microscope, the 3Di™ (Alpenglow Biosciences Inc.). Regions of interest (ROIs) with a volume of 0.5 mm3 were re-imaged at 0.33 µm/pixel. The volumetric imaging time was approximately 0.5 min per mm3 of tissue for each wavelength channel.

Images were processed using Alpenglow’s automated 3Dm processing pipeline, which performs stitching, registration, and fusion. The processed volumetric images were subsequently converted computationally to a standard H&E color format using Fiji and Aivia (Leica) software and visualized using Aivia software.

### 3D H&E reconstruction with CODA

#### Image registration

To reconstruct a 3D tissue volume from serial sections, foreground tissue in consecutive H&E images was first detected by mean-thresholding the Gaussian-filtered (R + B)/2 minus G channel difference, with any background pixel below the average magenta-minus-green difference set to white. The 10× images were then registered using CODA (40) as described in the original paper, with the code version last updated on Dec 13, 2023.

#### Semantic segmentation

7 of 36 consecutive sections were manually annotated by our clinical pathologist for training across 10 classes: germinal centers, macrophage, fibroinflammatory stroma, necrosis, background lymph node, blood vessel, fibrosis, fat, calcification, and fibrotic nodule, with fat also serving as the non-tissue background class. Using these annotations, the modified DeepLab V3 ResNet-50 model implemented in CODA was trained with n_train = 15, sxy = 700, annotation_whitespace = [2 2 2 2 2 2 2 2 2 2], add_whitespace_to = [8 8], nesting_order = [6 1 9 2 3 7 5 4 10 8], combine_classes = [1 2 3 4 5 6 7 8 9 10], and otherwise default parameters. The colored tissue block was reconstructed by stacking the registered, labeled masks along the z-axis.

### QUANTIFICATION AND STATISTICAL ANALYSIS

Details of the statistical tests conducted in the study are indicated in the Results section and Figure Legends and summarized as below:

1) For the comparisons of cell type percentages, only cells inside the annotated tumor bed areas were included in the calculation, except for uninvolved lymph node samples. The data is not normally distributed, so the Mann-Whitney U test was used to test for statistical significance between response groups.
2) For comparisons of TLS and GC behavior quantified with GCSCAN, only cells and structures within the annotated tumor bed were included. A TLS or follicle was scored as GC-positive when at least 15% of its cells were CD21⁺Ki67⁺ and located inside the approximated GC area. Comparisons were made between response groups or between combined response-treatment groups. Follicle size was summarized per sample by the sample mean, and the within-follicle GC ratio by the upper quartile, chosen to better capture the shifted GC distribution. Because the data were not normally distributed, the Mann-Whitney U test was used for significance.
3) For pairwise comparisons in analyses involving more than two groups, p-values were adjusted for multiple comparisons using the Holm-Bonferroni method (package statsmodels, version 0.14.1).

